# Engineering of Phytosterol-Producing Yeast Platforms for Functional Reconstitution of Downstream Biosynthetic Pathways

**DOI:** 10.1101/2020.06.08.139279

**Authors:** Shanhui Xu, Curtis Chen, Yanran Li

## Abstract

As essential structural molecules for plant plasma membranes, phytosterols are key intermediates for the synthesis of many downstream specialized metabolites of pharmaceutical or agricultural significance, such as brassinosteroids and withanolides. *Saccharomyces cerevisiae* has been widely used as an alternative producer for plant secondary metabolites. Establishment of heterologous sterol pathways in yeast, however, has been challenging due to either low efficiency or structural diversity, likely a result of crosstalk between the heterologous phytosterol and the endogenous ergosterol biosynthesis. For example, in this study, we engineered campesterol production in yeast using plant enzymes; although we were able to enhance the titer of campesterol to ~40mg/L by upregulating the mevalonate pathway, no conversion to downstream products was detected upon the introduction of downstream plant enzymes. Further investigations uncovered two interesting observations about sterol engineering in yeast. First, many heterologous sterols tend to be efficiently and intensively esterified in yeast, which drastically impedes the function of downstream enzymes. Second, yeast can overcome the growth deficiency caused by altered sterol metabolism through repeated culture. By employing metabolic engineering, strain evolution, fermentation engineering, and pathway reconstitution, we were able to establish a set of phytosterol-producing yeast strains with decent growth and titer of campesterol (~ 7mg/L), β-sitosterol (~2mg/L), 22-hydroxycampesterol (~1mg/L), and 22-hydroxycampest-4-en-3-one (~4mg/L). This work resolves the technical bottlenecks in phytosterol-derived pathway reconstitution in the backer’s yeast and opens up opportunities for efficient bioproduction and pathway elucidation of this group of phytochemicals.

## Introduction

Phytosterols are cholesterol-like sterols produced from plants that exhibit distinct side chains from cholesterol. Similar to cholesterol in humans and ergosterol in yeast, phytosterols are essential membrane components that regulate membrane fluidity and permeability in plants^1, 2^. Campesterol, **1** and β-sitosterol, **2** are the dominant sterols in plants^3^, and their biosynthesis has been relatively well elucidated^4–6^. In addition to their essential role in membrane maintenance, phytosterols exhibit a broad spectrum of promising bioactivities. Phytosterols are well-documented for their cholesterol-lowering effect^3, 7^ and have been listed as a dietary factor that may affect cardiovascular disease risk by the American Heart Association^8^. Phytosterols have also been reported to be associated with reduction in cancer risks^9, 10^, anti-oxidant^11, 12^ and anti-osteoarthritic properties^3, 13^. In addition, phytosterols serve as the synthetic precursor of downstream phytosteroids, such as the important group of plant hormones brassinosteroids (BRs), and pharmaceutically intriguing withanolides.

BRs are one important group of plant hormones involved in a number of physiological functions and behaviors^14, 15^, such as cell division, vascular differentiation, and pollen elongation. BRs exhibit extremely low abundance in nature (e.g. 10-100mg/kg in pollen, 10-100ng/kg in shoots and leaves), which makes them difficult to isolate, detect, and utilize. In addition, BRs are of broad pharmaceutical interests with anticancer, antiviral, and anabolic activities^16^. Brassinolide is the best studied and the most active brassinosteroid^17^, and is synthesized from campesterol^18^. Most BRs are synthesized along the brassinolide biosynthetic pathway from campesterol, while a small number of others were proposed to be synthesized from β-sitosterol^19–21^. Enzymes involved in the early and final stage of brassinolide biosynthesis have been identified, while the ones catalyzing the reactions between these enzymes remain unclear (Figure 1). In addition, how the enzymes catalyzing early steps are arranged to direct the synthesis towards brassinolide is unknown. One of the hindrances to clearly elucidate the biosynthesis is the lack of intermediates required to examine substrate preference of the enzymes involved and reconstitute the biosynthesis *ex situ*. Ergosterol, **3**, biosynthesis and phytosterol biosynthesis diverge from as early as squalene; while campesterol biosynthesis comes across ergosterol biosynthesis at the synthesis of episterol, **4**, which is converted to ergosterol or campesterol in fungi or plant, respectively (Figure 1). The bioproduction of campesterol has been achieved in *Saccharomyces cerevisiae*^2^ and *Yarrowia lipolytica*^23^, through replacing the endogenous C-22 sterol desaturase (ERG5) gene with a heterologous 7-dehydrocholesterol reductase (DHCR7) encoding gene. In plant, campesterol is synthesized from **4** by the function of Δ^7^-sterol-C5-desaturase (DWF7), 7-dehydrocholesterol reductase (DWF5) and Δ^24^-sterol reductase (DWF1)^24^. On the other hand, the bioproduction of β-sitosterol has not been established in heterologous microbial hosts, which is likely due to the lack of substrate for the sterol-C24-methyltransferase SMT2.

The recent development of plant pathway reconstitution in heterologous hosts has highlighted the potential of using microbial hosts (e.g. *S. cerevisiae*) for pathway elucidation and enzyme characterization in natural product biosynthesis. *S. cerevisiae* has been utilized for enzyme characterization in phytosterol and derivative biosynthesis. For instance, the desmosterol-producing yeast and 24-methylenecholesterol-producing yeast constructed through expressing DWF5 into *erg6* and *erg4/erg5* inactivated strains, respectively, have been utilized to characterize sterol Δ^24(25)^ reductase involved in cholesterol biosynthesis^25^, novel sterol Δ^24(28)^ reductase involved in campesterol production^26^, and Δ^24^ isomerase putatively used in withanolide biosynthesis^27^. The cholesterol-producing yeast constructed through expressing 24-dehydrocholesterol reductase (DHCR24) and DHCR7 in the *erg5/erg6* inactivated yeast strain has been utilized to confirm the function of CYP90B27 in synthesizing 22(R)-hydroxycholesterol from cholesterol^28^. The same cholesterol-producing yeast strain has also been utilized to reconstruct diosgenin biosynthesis with the expression of *Pp*CYP90G4/*Pp*CYP94D108 or *Tf*CYP90B50/*Tf*CYP82J17. However, no β-sitosterol or campesterol-derived compounds have been successfully synthesized in yeast.

**Figure 1.**
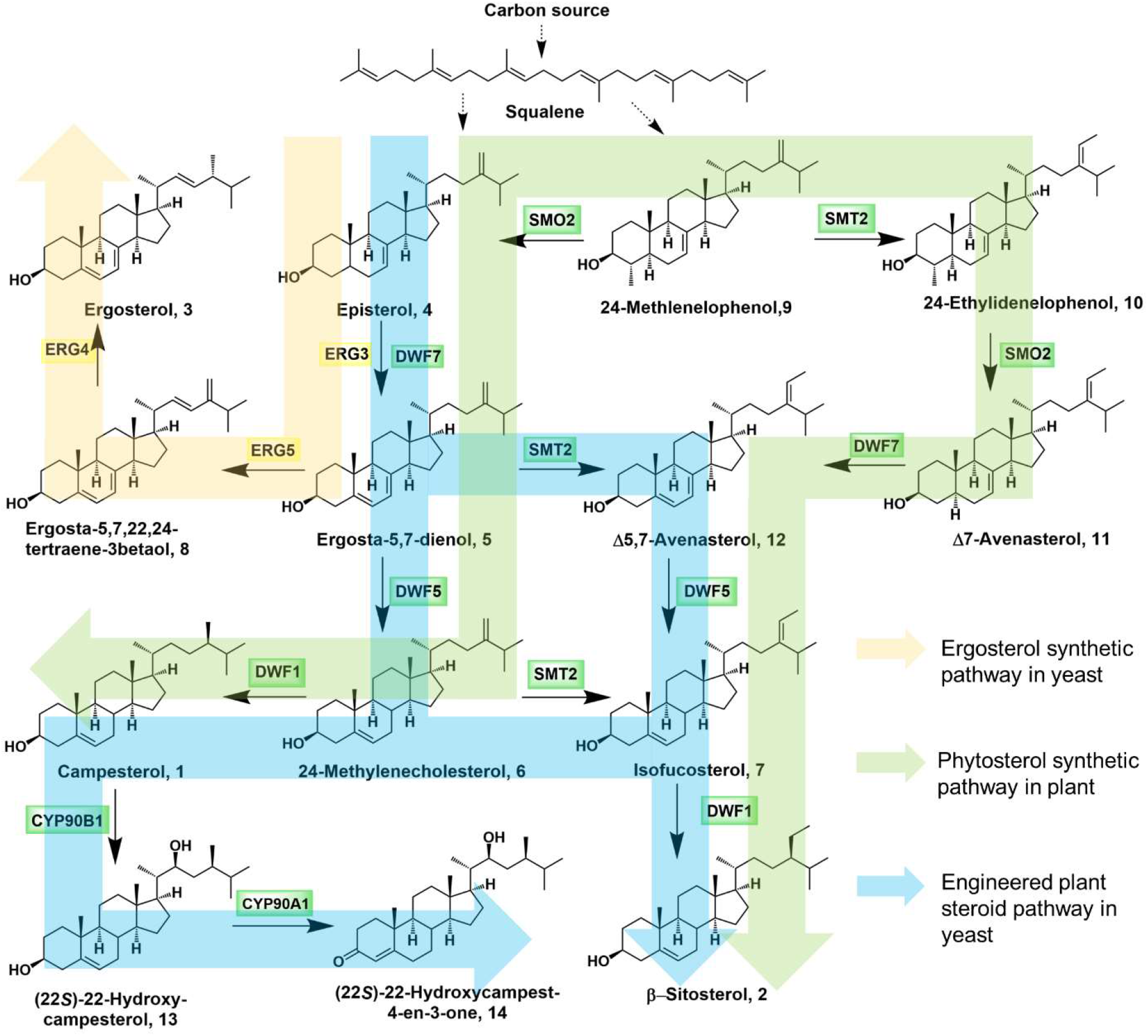
Proposed biosynthetic pathway of phytosterol and (22S)-22-Hydroxycampest-4-en-3-one, **14** in plant and yeast. The yellow arrow represents the native ergosterol, **3**, biosynthetic pathway in yeast; the green arrow represents the indigenous pathway of β-sitosterol, **2**; the green arrow represents the reconstituted biosynthetic pathway from episterol, **4** to **14**.

Here, we resolved several bottlenecks in phytosteroid-related pathway reconstitution in *S. cerevisiae* and demonstrated the establishment of a yeast-based biosynthetic platform that enabled an efficient reconstitution of the early-stage brassinolide biosynthetic pathway towards the synthesis of 22-hydroxycampest-4-en-3-one, **14**. The biosynthesis of campesterol from **4** was established in yeast using plant enzymes. Through upregulating the mevalonate pathway, campesterol titer was enhanced to ~40mg/L in yeast, from ~3mg/L. However, campesterol and pathway intermediates were intensively esterified and thereby inaccessible by subsequent plant enzymes. Removal of sterol acyltransferase genes and *erg4*, though led to significant growth burden, enabled the synthesis of β-sitosterol in yeast by the function and substrate promiscuity of SMT2. Surprisingly, the engineered yeast strain of altered sterol composition efficiently adapted itself to overcome the growth deficiency through repeated culture. Genome sequencing analysis indicates a number of mutations from the original strain that may explain this adaptation although the exact mechanism remains to be experimentally confirmed. Furthermore, we were able to establish and optimize the biosynthesis of **14** in yeast, which not only enabled us to confirm the reaction sequence of the two cytochrome P450s, but also provided an efficient **14**-producing strain for the reconstitution and elucidation of the downstream pathway towards BR synthesis.

## Results

### Construction and optimization of *de novo* campesterol production using plant enzymes in yeast

Campesterol, **1** and ergosterol, **3** are both biosynthesized from episterol, **4**, and this enables the redirection of yeast ergosterol synthetic pathway to the synthesis of campesterol. Plants use DWF7, DWF5 and DWF1 to convert **4** to campesterol (Figure 1). DWF7, the ortholog of ERG3, catalyzes the conversion of **4** to ergostra-5,7-sterol, **5**. **5** is then converted to 24-methylenecholesterol, **6** through C7-C8 enol reduction by DWF5, followed by C24-C28 enol reduction catalyzed by DWF1 to afford the formation of campesterol. Since DWF7 and ERG3 are functional exchangeable, heterologous expression of DWF5 and DWF1 should lead to the synthesis of campesterol in yeast. The *dwf5* and *dwf1* genes from *Arabidopsis thaliana* were introduced into yeast with low-copy plasmids and regulated by the yeast constitutive *GPD* promoter. The engineered yeast was grown in synthetic defined medium (SDM) for 3 days at 30°C. Since sterols were mainly present in yeast cells^29^, cells were collected and saponified with 60% (w/v) KOH using the standard protocol^30^. The extracted samples were analyzed with liquid chromatography-mass spectrometry (LC-MS) and the synthesis of campesterol in yeast was confirmed by comparing with the authentic campesterol standard (Figure 2A). Further introduction of *dwf7* to the *dwf5/dwf1*-expressing strain resulted in significant enhancement in campesterol production (Figure 2A) at ~2mg/L, and minor effects on yeast growth (Figure S1). As a comparison, introduction of an extra copy of *erg3*, the *dwf7* ortholog in yeast, also resulted in the similar boost of campesterol production (Figure S2A), which suggests that ERG3 or DWF7 might catalyze a rate-limiting step in campesterol biosynthesis in yeast. Additionally, the introduction of *dwf7, dwf5* and *dwf1* genes into yeast strain greatly impacted the synthesis of ergosterol, which was barely detected from the engineered campesterol-producing yeast strains (Figure S2B). The *dwf7, dwf5 and dwf1* genes were then integrated onto the genome to afford a stable campesterol-producing strain for further engineering (YYL55).

**Figure 2.**
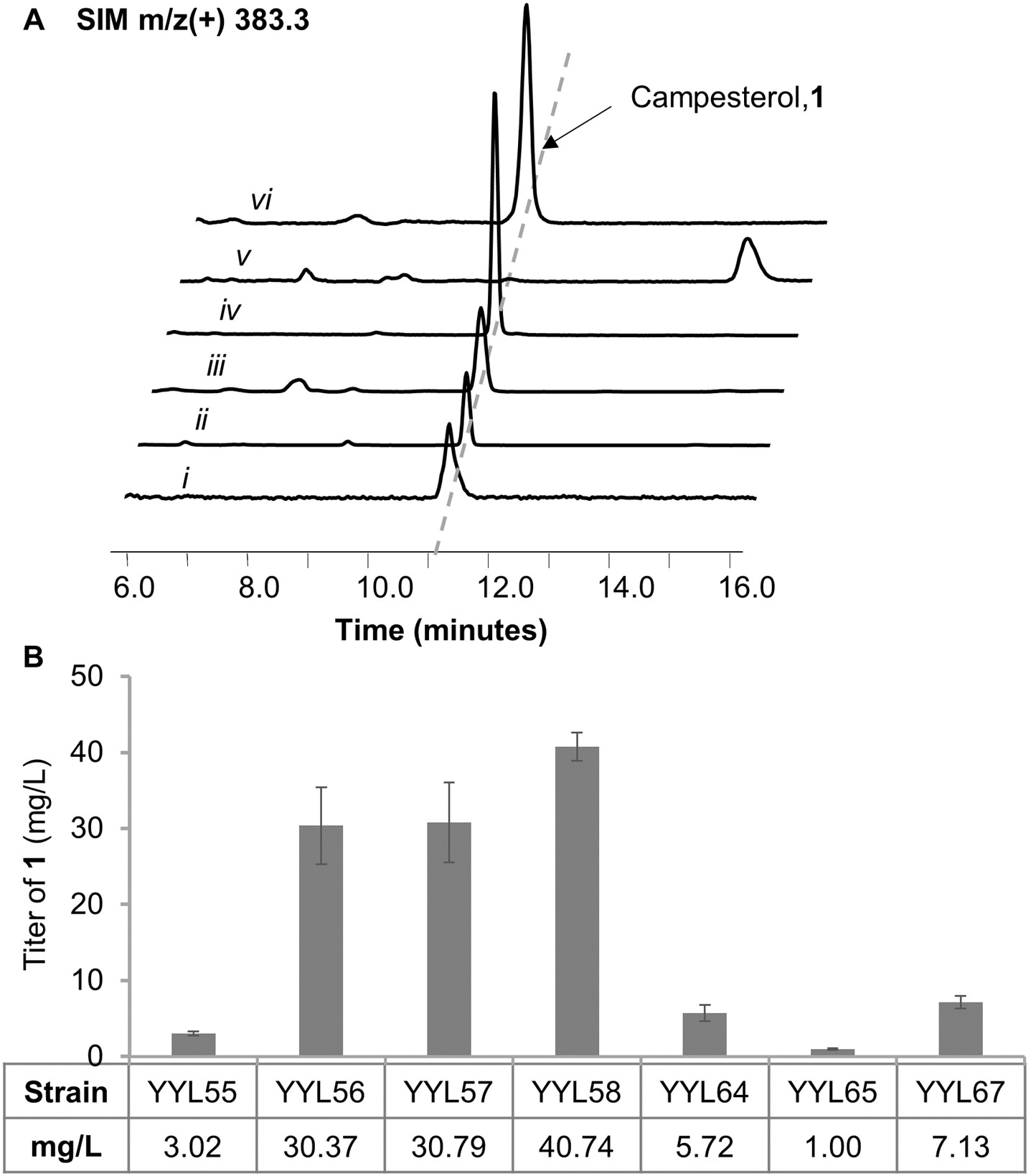
Biosynthesis of campesterol in yeast. **(A)** Selected ion monitoring (SIM) extracted ion chromatogram (EIC) using campesterol’s characteristic m/z^+^ signal (MW=400.69Da, [C_28_H_49_O-H_2_O]^+^=[C_28_H_47_]^+^=383.4) of i) campesterol standard, ii) yeast strain harboring DWF5, iii) yeast strain harboring DWF5 and DWF1, iv) yeast strain harboring DWF5, DWF1 and DWF7 (YYL56), v) YYL56 without KOH saponification, vi) YYL64 without KOH saponification. **(B)** Campesterol production from yeast strains harboring DWF7/DWF5/DWF1 with enhanced expression level of mevalonate pathway, *are1/are2* inactivation and *erg4* inactivation. The campesterol-producing yeast strains were cultured in synthetic complete SDM medium supplemented with 2% (w/v) glucose at 30°C for 72 hours. All traces are representative of at least three biological replicates for each engineered yeast strain. Bars represent mean values of three biological replicates, and the error bars represent the standard deviation of the replicates.

To increase the campesterol production to a level sufficient for the efficient reconstitution of downstream phytosteroid pathways, the endogenous mevalonate (MVA) pathway was upregulated in yeast to increase the synthesis of squalene, the precursor of sterols. Based on a previously established pathway^31^: *erg7, erg10, erg12, erg13, erg19, erg20* and *idi1* were overexpressed by strong, constitutive promoters in the yeast genome. Earlier reports also indicate that overexpressing the truncated 3-hydroxy-3-methylglutaryl-coenzyme A reductase (tHMG1) and the sterol regulatory element binding protein UPC2 could increase the production of squanlene^31, 32^. Thus, three squalene-upregulating modules (module I: *thmg1, erg10, erg12*, and *erg13*; module II: *erg8, erg19, idi1*, and *thmg1*; module III: *erg7, erg20, upc2*, and *thmg1*) were integrated sequentially into *ybl059w, ybr197c*, and *ymr206w* locus of YYL55 to afford YYL56, YYL57, and YYL58, respectively (Table S1). Consistent with previous studies and as expected, the overexpression of genes on MVA pathway improved campesterol production. Out of all the strains, YYL58 exhibited the highest production of campesterol at 40.72 mg/L, a 13-fold increase from the original campesterol-producing YYL55. However, upregulating squalene synthesis also led to a deleterious effect on the growth of yeast: compared with wild-type strain, YYL56, YYL57 and YYL58 all exhibited slightly longer lag phases and slower growth rates (Figure S1).

### Construction and characterization of β-sitosterol biosynthesis in yeast

Episterol, **4** is an important precursor of ergosterol and campesterol in fungi and plants, respectively. In plants, **4** is proposed to be synthesized from 24-methylenelophenol, **9**, by the function of the ERG25 ortholog sterol 4α-methyl oxidase SMO2^33^. **9** is proposed to be converted to 24-ethylidenelophenol, **10**, with the function of the sterol-C24-methyltransferase SMT2^34^, and SMO2, DWF7, DWF5, and DWF1 then function to convert **10** to β-sitosterol, **2** (Figure 1). Due to the structural similarity between **9** and **4**, and the fact that sterol pathway in plants is highly cross talked, it is likely that SMT2 could function on C24 methylene sterols, such as **4**, **5** or **6**. Therefore, SMT2 was cloned from *A. thaliana* cDNA library and expressed under a strong constitutive *GPD* promoter in YYL58. However, no β-sitosterol or any C24’-methylated products were detected (Figure 3). LC-MS analysis of the SMT2-expressing YYL58 indicated that **6** was accumulated in saponified extracts of the cell pellets, while **4** and **5** were barely detected, which is likely due to the efficient conversion of **4** and **5** to Ergosta-5,7,22,24-tertraene-3betaol, **8**, due to the high efficiency of ERG3 and ERG5 enzymes. Normally, surplus or foreign sterols are esterified and stored in lipid droplets in yeast to reduce the toxicity^35^. This mechanism of regulating steroid synthesis by sterol esterification is also found in plants. Previous investigations found that changing the expression level of a putative acyl transferase BAT1 (BR-related acyltransferase 1) may influence BR levels in plants: overexpression of BAT1 in *A. thaliana* leads to BR-deficient phenotypes, which can be rescued by BR feeding^36^. Thus, we hypothesize that **6** may be esterified in yeast and became inaccessible to SMT2. To confirm this possibility, YYL58 cell pellets were extracted without saponification. Interestingly, compared with the saponified samples, **6** and campesterol were barely detected in the unsaponified extracts of YYL58, indicating highly efficient esterification of these two phytosterols in yeast (Figure S3). Complete esterification of campesterol and **6** was found in all the other campesterol producing strains as well.

To enhance the availability of free phytosterols in yeast, two acyltransferases responsible for sterol esterification were selected to be deleted from YYL58. Previous investigations indicate that two acyl-CoA sterol acyltransferases ARE1 and ARE2 are responsible for the esterification of sterols in *S. cerevisiae*^31, 38^. Inactivation of *are1* and *are2* demolished the synthesis of sterol ester and enhanced free sterols levels^37, 38^. However, inactivation of *are1* and *are2* was lethal to YYL58, likely due to the accumulation of free phytosterols, isopentenyl pyrophosphate (IPP), or dimethylallyl pyrophosphate (DMAPP)^39^. YYL56, the campesterol-producing strain exhibited more robust growth than YYL58 (Figure S1) but with a similar level of campesterol production, was selected for *are1* and *are2* inactivation to generate YYL64.

**Figure 3.**
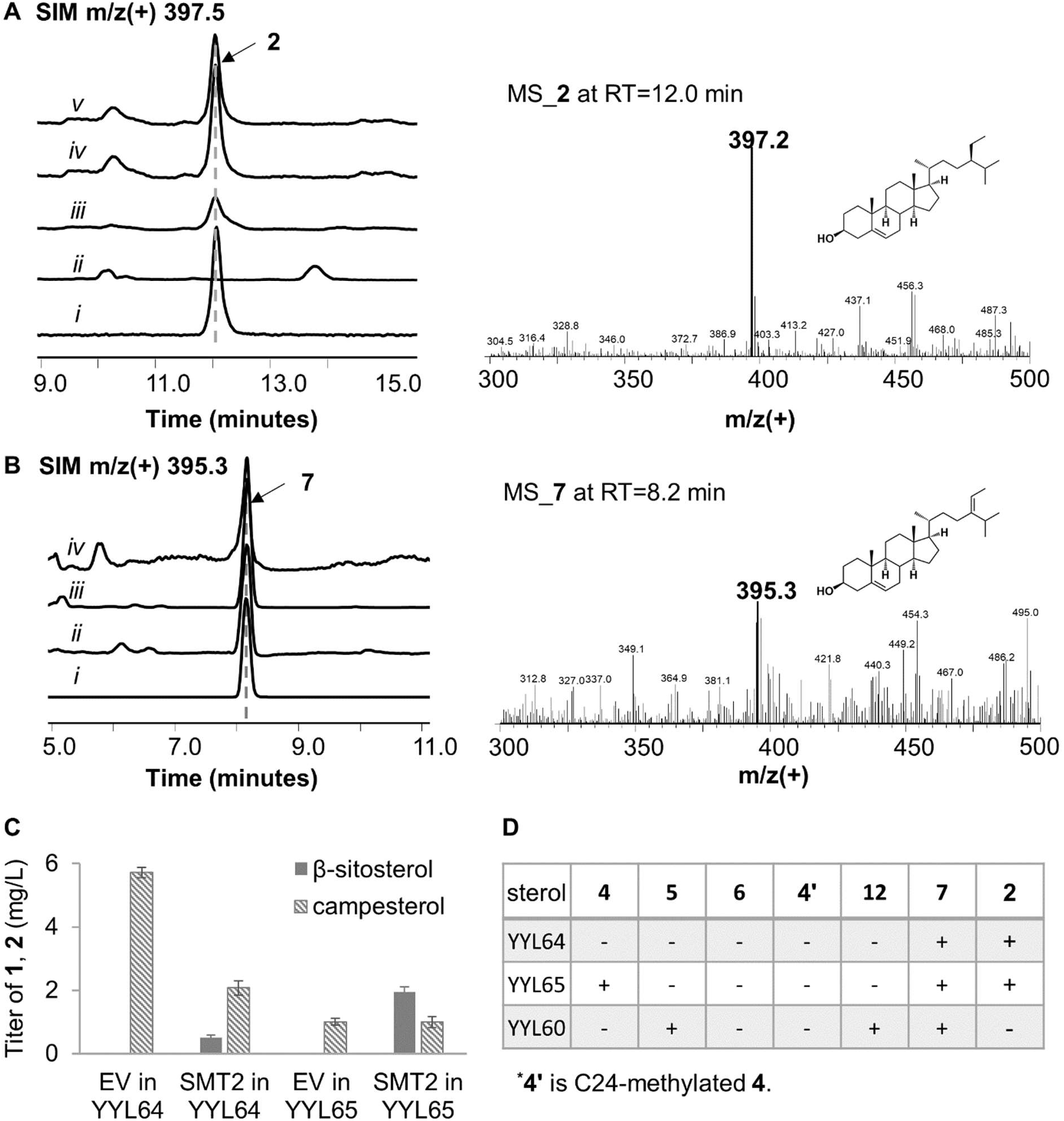
Biosynthesis of β-sitosterol in yeast. **(A)** Selected ion monitoring SIM (SIM) extracted ion chromatogram (EIC) using β-sitosterol’s characteristic m/z^+^ signal (MW=414.71Da, [C_29_H_51_O-H_2_O]^+^=[C_29_H_49_]^+^=397.4) of i) β-sitosterol standard, ii) YYL56 harboring SMT2, iii) YYL64 harboring SMT2, iv) YYL65 harboring SMT2, v) YYL67 harboring SMT2. **(B)** EIC SIM using isofucosterol, **7**, characteristic m/z^+^ signal (MW=412.70Da, [C_29_H_49_O-H_2_O]^+^=[C_29_H_47_]^+^ =395.3) of i) isofucosterol standard, ii) YYL64 expressing SMT2, iii) YYL65 expressing SMT2, iv) YYL67 expressing SMT2. **(C)** Campesterol and β-sitosterol production in YYL64 and YYL65 expressing SMT2. **(D)** Sterol composition of YYL64, YYL65 and YYL60 in the presence of SMT2. YYL64 and YYL65 were extracted without saponification; YYL60 was extracted with saponification method. All traces are representative of at least three biological replicates for each engineered yeast strain. Bars represent mean values of three biological replicates, and the error bars represent the standard deviation of the replicates.

Compared to YYL56, YYL64 synthesized campesterol and **6** as free sterols, with a decreased campesterol titer at 5.72 mg/L (Figure 2B). However, the introduction of SMT2 in YYL64 resulted in β-sitosterol production although at only ~0.5mg/L and with a cost of campesterol production dropped around 60%, which was synthesized by the same set of enzymes except for SMT2 (Figure 1, 3A, 3C). The significantly reduced production of β-sitosterol compared to campesterol in YYL64 expressing SMT2 indicates that the synthesis of β-sitosterol is likely due to the promiscuity of SMT2, which is consistent with previous investigations that the synthesis of β-sitosterol and campesterol branches from 23-methlenelophenol, **9**, the putative native substrate of SMT2. In SMT2-expressing YYL64, isofucosterol, **7** was detected while the methylated products of **4** and **5** were not, which indicates that **6** can be recognized by SMT2 due to enzyme promiscuity (Figure 1, 3B, 3D). The inactivation of *are1* and *are2* in YYL56 also led to growth defects, altered the sterol composition, and diminished total sterol production in YYL64 (Figure S2). Since the inactivation of *are1* and *are2* in wild type strain (YYL63, Table S1) had a minimal effect on yeast viability (Figure S1), the growth defect in YYL56 is likely due to the accumulation of foreign sterols or squalene. The weakened growth of YYL64 in comparison with YYL56 also implies the importance of esterification in reducing the toxicity of free foreign sterols in yeast.

ERG4 and DWF1 are both Δ24-sterol reductases that function on the C24 double carbon bond. When functioning on 24-methylenecholesterol, DWF1 catalyzes the formation of both 24(S)- and 24(R)-campesterol; while only 24(R)-β-sitosterol is synthesized in plants^26, 40^. On the other side, ERG4 has only been reported to synthesize 24(*S*)-methyl sterol, ergosterol, but not C24-ethyl sterols such as β-sitosterol^41^. Consistent with previous results that campesterol could be synthesized from DHCR7-expressing yeast^22, 23^, expressing DWF5 in wild type yeast led to the synthesis of campesterol; while no campesterol was detected when DWF5 was introduced to the *erg4Δ* mutant strain(Figure 2B, S4). These facts imply that while DWF1 can function on both **6** and **7**, ERG4 may only function on **6**. Thus, ERG4 likely competes with SMT2 in the conversion of **6** in SMT2-expressing YYL64. To improve the synthesis of β-sitosterol, *erg4* was then deleted from YYL64 to generate YYL65. Inactivation of *erg4* resulted in growth deficiency, such as the hypersensitivity to certain drugs and impaired mating process^41^. Here, inactivation of *erg4* resulted in dramatically reduced campesterol synthesis (~1mg/L) (Figure 2) and a significant growth burden on YYL65, which exhibits an extended lag phase and largely decreased cell density at the stationary phase (Figure 4, Figure S1). However, the expression of SMT2 in YYL65 resulted in an enhanced production of β-sitosterol at 2mg/L, an approximately 4-fold increase compared to YYL64 (Figure 3A, 3C). Different from what we observed in YYL64, introduction of SMT2 in YYL65 did not affect campesterol production (Figure 3B); although **6** was not detected, **7** was observed in the SMT2-expressing YYL65 (Figure 3D). The decreased cell density with a higher β-sitosterol production, compared to YYL64, agreed with the possibility that ERG4 may compete with SMT2 in β-sitosterol synthesis, and **6** is a major substrate of SMT2 in YYL64 and YYL65.

**Figure 4.**
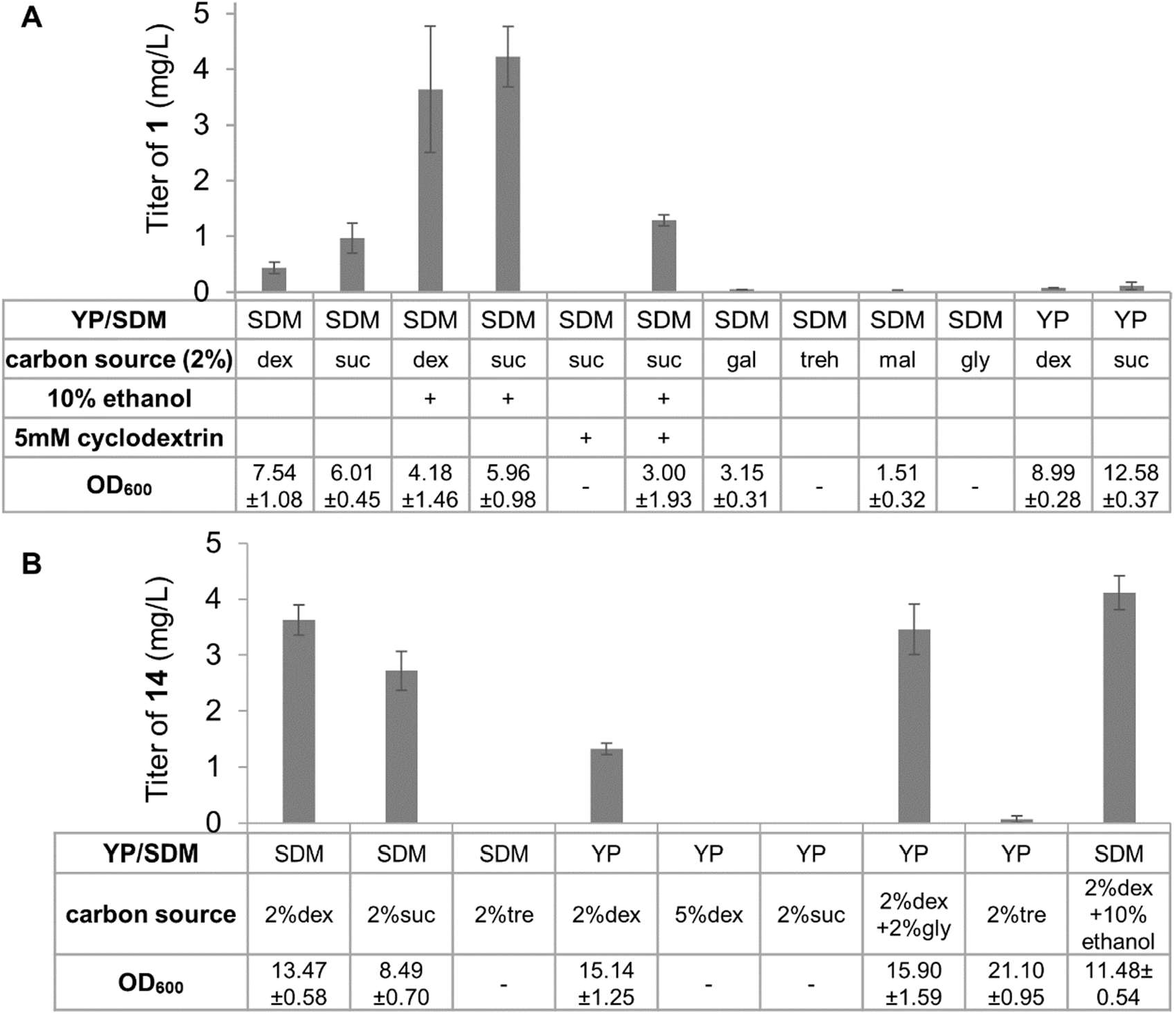
Medium optimization of **(A)** YYL65 and **(B)** YYL69. SDM represents as synthetic defined minimal medium with 1.7g/L yeast nitrogen bases, corresponding dropout and 5g/L ammonium sulfate, pH 5.8. YP represents complex medium with 10 g/L yeast extract, 20 g/L peptone and 80mg/L adenine. 2% (w/v) of various carbon sources were used: dex (D-glucose), suc (sucrose), gal (galactose), treh (trehalose), mal (malose) and gly (glycerol). Bars represent mean values of three biological replicates, and the error bars represent the standard deviation of the replicates.

In addition, to identify the possible substrates of SMT2 in addition to **6** in yeast, the methylated products of **4** and **5** were examined in **4** and **5** accumulating strains. Methylated episterol, **4’**, was not detected in YYL65, which accumulated **4**. It suggested that **4** cannot be converted by SMT2 in yeast (Figure 3D). We also introduced SMT2 to the **5**-accumulating strain YYL60 (Table S1), and observed the synthesis of a compound with m/z^+^=393.4 that agrees with **12** ([C_29_H_47_O-H_2_O]^+^=[C_29_H_45_]^+^=393.4, Figure S5), indicating that SMT2 can also function on **5**. Different from **6** and campesterol, we were able to detect free **5** in YYL60 (Figure S3C). Taken together, these results suggest that **5** and **6** maybe the substrates of SMT2 in YYL64 and YYL65 that route the synthesis towards β-sitosterol upon the introduction of SMT2 (Figure 1).

### Optimization of phytosterol production in yeast

The successful construction of β-sitosterol production enabled us to identify the obstacles in metabolic engineering of phytosterol synthesis in yeast. The cross talk between the endogenous sterol metabolism and the heterologous sterol pathway often make the substrates not readily accessible for the downstream heterologous enzymes. Thus, disruption of sterol metabolism may be necessary for the successful reconstitution of phytosterol biosynthesis, but this also likely introduce growth deficiencies, such as in the case of YYL65. Due to the growth deficiency introduced by the *are1/are2* and *erg4* inactivation and the relatively low titer of campesterol, functional reconstitution of more heterologous enzymes in YYL65 became unlikely.

To improve phytosterol production and alleviate the growth stress, we optimized the medium for YYL65. Different carbon sources were used to improve the growth condition and enhance campesterol production (Figure 4A). We also examined the effects of methyl-β-cyclodextrin considering that it may alleviate the sterol stress in YYL65 by transporting deleterious sterols out of yeast^42^. The effect of supplementing 2% (w/v) sucrose with different amounts of ethanol (2%, 5%, and 10%, v/v) was examined as well (Figure S6). Interestingly, supplementing 10% (v/v) ethanol to SDM with 2% (w/v) sucrose or glucose led to up to 10-fold increase in campesterol production but also extended the lag phase of yeast growth. The enhanced production of campesterol is consistent with previous reports of yeast production of amorphadiene, which is likely due to the fact that ethanol enhanced the supply of acetyl-CoA^32^. In addition, a high concentration of ethanol was reported to trigger an increased ergosterol synthesis in order to protect damaged plasma membrane^43^. Thus, the addition of ethanol may upregulate sterol synthesis and result in enhanced supply of **4** and **5**, substrates of campesterol. However, despite the elevated campesterol titer in the presence of ethanol, we did not observe obvious alleviation of growth stress in the medium conditions tested.

Meanwhile, we noticed that re-streaking YYL65 on agar plates resulted in two different morphologies of yeast colonies – some exhibited a much larger size than the remaining majority, indicating that some yeast cells may have mutated to circumvent the growth burden caused by the altered sterol composition. Based on this interesting observation, we conducted adaptive evolution on YYL65 to screen for derivatives with enhanced growth. For inoculum preparation, YYL65 was cultivated at 250 rpm and 30°C in 2mL SDM till it reached the stationary phase (OD600 ~6.0). An appropriate volume of this culture was used to inoculate 2mL fresh SDM with an initial OD600 of 0.1. Once the culture reached early stationary phase, serial dilutions were repeated. In this procedure, the time interval that YYL65 took to reach the stationary phase became shorter after each cycle, indicating enhanced growth. The culture after 10 cycles were plated on agar plate and 10 individual colonies were further analyzed by culturing in SDM with and without 10% (v/v) ethanol. Campesterol titer was then quantified by LC-MS. Excitingly, all 10 colonies grew more robustly than the original YYL65 and the production of campesterol was all boosted. Seven colonies synthesized a higher amount of campesterol in the presence of 10% (v/v) ethanol and one colony didn’t survive in the high concentration of ethanol (Figure 5A). Among them, YYL65-5 had the highest production of campesterol in the presence of ethanol, while YYL65-1 and YYL65-2 exhibited comparable level of campesterol titer without the presence of ethanol (Figure 5A). YYL65-1, with both an enhanced growth pattern and a higher campesterol production in normal medium condition (campesterol titer at ~ 7.13mg/L, Figure 2B, Figure 5B), was annotated as YYL67 and selected for the further pathway reconstitution.

**Figure 5.**
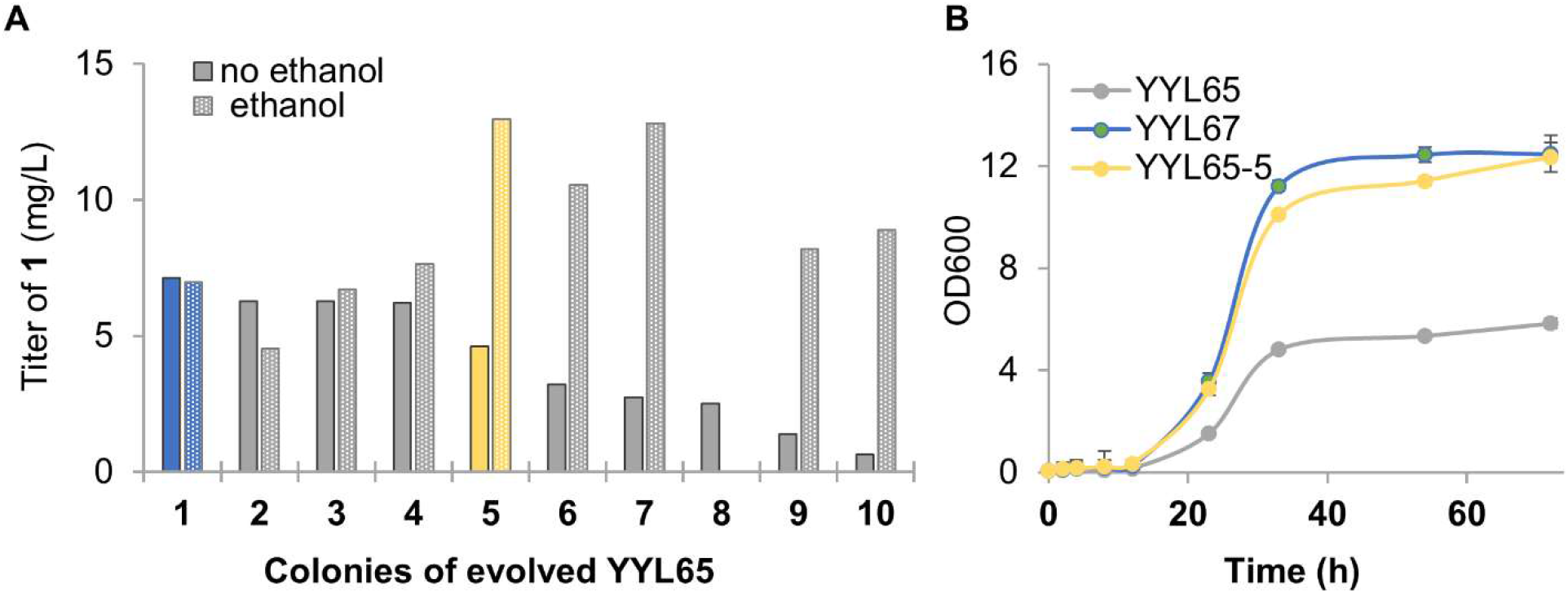
Adaptive evolution of YYL65. **(A)** Campesterol production of YYL65 and the ten mutants picked from the 10th generation of adaptive evolution in synthetic complete SDM supplemented with 2% sucrose, with and without 10% (v/v) ethanol. **(B)** Growth curve of YYL65, YYL65-7, and YYL67. The yeast strains were cultured in synthetic complete SDM medium at 30°C for 72 hours.

We next analyzed the genome sequences of YYL65, YYL65-5, and YYL67 to understand the possible mechanism underlying the enhanced growth and campesterol production observed in YYL65-51 and YYL67. Paired-end libraries (150-bp) were generated by NGS platforms and sequenced to an average depth of over 50x for each strain. The high coverage enabled reliable genome-wide analysis of insertion and deletion (InDel) and single-nucleotide polymorphisms (SNPs) (Figure S7, Table S2, S3, S4). Compared to the parental strain, YYL65, YYL67 and YYL65-5 carry 35 and 41 InDels in gene-coding regions (Table S3), respectively. 22 of these InDels are present in both YYL67 and YYL65-5 (Table S3); many of these genes are related to transcription regulation and growth stress response (Table S2B). We are particularly interested in *ASG1* and *SWI1*. ASG1 is an activator of stress response genes and the null mutants of *asg1* have a respiratory deficiency. a mutation with seven asparagine (N) inserted at C terminus N-enriched region is exclusively present in the ASG1 protein of YYL67. SWI1 is the subunit of the SWI/SNF chromatin remodeling complex and is required for the transcription of many genes. A mutated *SWI1* gene is present in both evolved strains but not in the parental strain YYL65. In addition, YYL65-5 and YYL67 have around 840 SNPs compared to the parental strain. Approximately 300 SNPs occur within open reading frames, which affect 60 and 65 genes in YYL67 and YYL65-5, respectively (Table S2). Interestingly, 49 genes affected by SNPs can be found in both evolved strains, making them good candidates that may contribute to the altered phenotypes. We noted that genes related to flocculation (*flo*1, *flo*5, *flo*9) were found in both SNP and InDel analysis. Flocculation is a process where yeast aggregates together to form a multicellular mass, which has been found to be correlated with stress tolerance development in *S. cerevisiea*^44, 45^. In addition, mutations were also observed in genes related to the multivesicular vesicle body sorting pathway (*cos*7, *cos*8, *cos*12) in YYL65-5 and YYL67, mainly in YYL65-5. The consistency of mutations in genes of related functions implies the importance of these genes in adapted yeast strains to accommodate the growth burden caused by altered sterol compositions. Although further experimental confirmations, and transcriptomic and metabolomic analyses are needed to understand the exact mechanism, the evolution of YYL65-7 and YYL67 from YYL65 highlights the ability of *S. cerevisiae* to adjust itself to the growth burden caused by altered sterol composition and supports the potential of using yeast for the bioproduction of phytosterols and their derivatives.

### Synthesis of 22-hydroxycampest-4-en-3-one in yeast

With the phytosterol-producing strain engineered to overcome the growth deficiency and produce phytosterols accessible for downstream enzymes, YYL67 can be utilized as a phytosterol-producing platform to reconstitute and elucidate the downstream biosynthetic pathway. Since the production of YYL67 in glucose and sucrose was similar (Figure S6), we switched to normal SDM supplemented with 2% glucose for the subsequent YYL67-related engineering and culturing. To explore this possibility, we attempted to produce BRs using YYL67. BRs are a group of universal hormones derived from campesterol and play essential roles in higher plants^14, 15^. They are involved in plant growth regulation, developmental processes, and responses to abiotic or biotic stress^14, 15, 46, 47^ For example, exogenous application of BR on potatoes have led to enhanced productivity and starch content^14^. The ability to boost crop yield makes BRs important biostimulants in agriculture. In addition, BRs also show promising antiviral, anticancer, and antitumor activities^20^. However, the low abundance of BRs in nature and the structural complexity make these molecules very difficult and expensive to obtain through direction isolation or chemical synthesis, and the high price and low accessibility of natural BRs of high purity and the lack of pathway intermediates are largely impeding the investigation and development of BRs for agricultural and pharmaceutical applications. Reconstituting BR pathways in microbial organism is a promising alternative approach to produce various BRs in high purity and economically. However, the biosynthetic pathways of BRs are not fully elucidated, with at least two enzymes missing and the sequence of the biosynthetic enzymes elusive. The construction of YYL67 enabled us to establish BR biosynthetic pathways in yeast, which may serve as a suitable platform for future BR production.

CYP90B1 was demonstrated as the first and rate-limiting step in BR biosynthesis^48^. Expression of CYP90B1 under the regulation of *GPD* promoter in YYL67, which contains NADPH–cytochrome P450 oxidoreductase ATR1, led to an efficient conversion of campesterol to (22 *S*)-22-hydroxy-campesterol, **13** (m/z^+^ = 399.3, Figure 6A, 6B), which is consistent with the previous *in vitro* biochemical characterization of CYP90B1^48^. The LC-MS/MS analysis further confirmed that the hydroxyl group was add at the C22 position (Figure 6B, S8). We also introduced CYP90B1 to YYL58 and YYL64, but the synthesis of **13** was either undetected or inconclusive (Figure 6A). We were unable to express CYP90B1 in YYL65 due to significant growth deficiency. The failed functional reconstitution of CYP90B1 in YYL58, YYL64, and YYL65 again highlights the importance of inactivating sterol acyltransferase genes and *erg4* from YYL56, and the directed evolution of YYL65 for the efficient construction of plant metabolite biosynthesis downstream of simple phytosterols (e.g. **1**, **2**, **6**) in *S. cerevisiae*.

**Figure 6.**
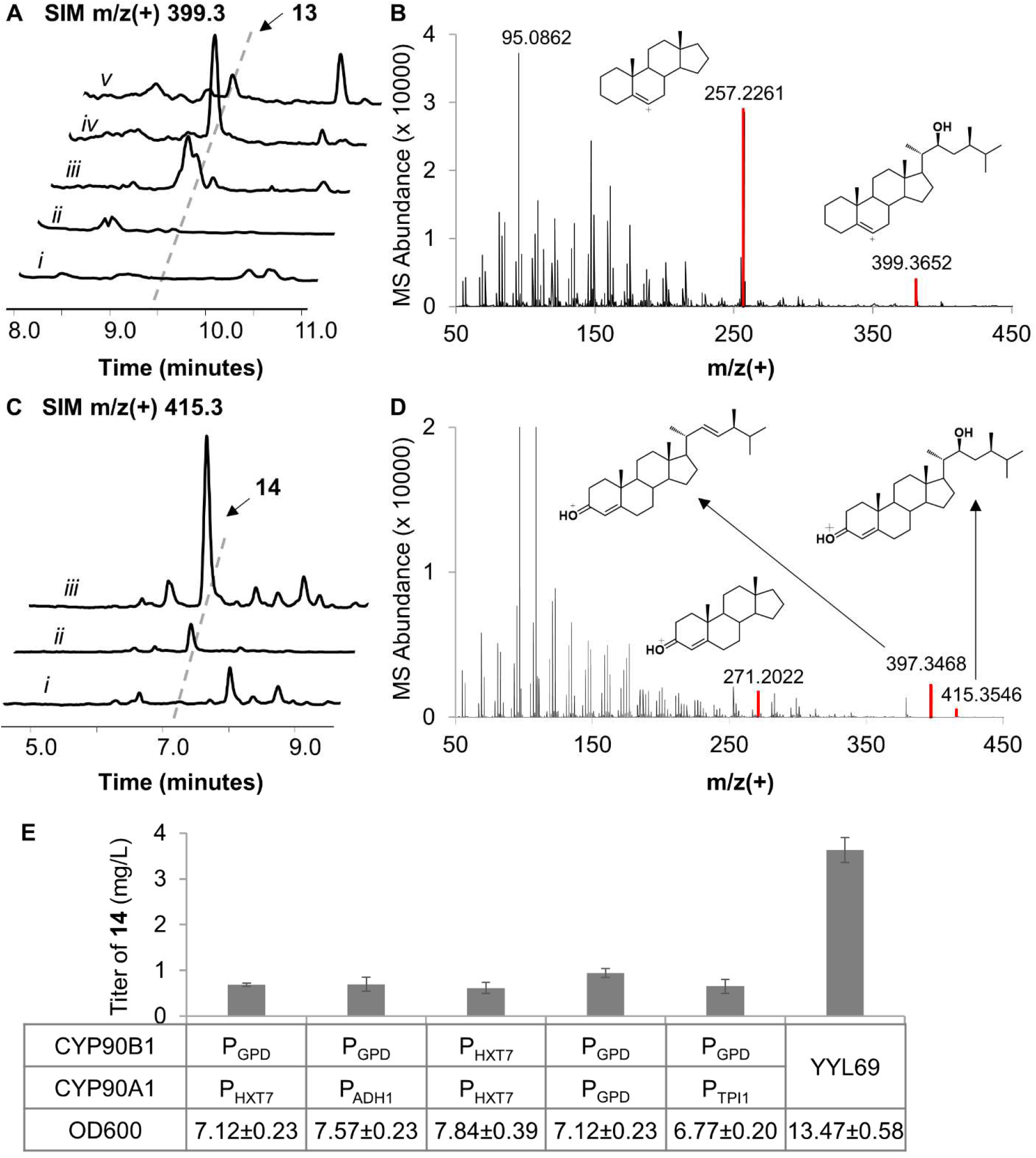
Biosynthesis of (22S)-22-Hydroxycampest-4-en-3-one, **14**. **(A)** SIM EIC using (22S)-22-Hydroxy-campesterol, **13**’s characteristic m/z^+^ signal (MW=416.69Da, [C_28_H_49_O_2_-H_2_O]^+^ [C_28_H_47_O]^+^=399.3) of i) YYL67, ii) YYL56 expressing CYP90B1, iii) YYL64 expressing CYP90B1, iv) YYL67 expressing CYP90B1, v) YYL67 expressing CYP90B1 and CYP90A1. **(B)** High resolution mass spectrum of **13**. **(C)** SIM EIC using (22S)-22-Hydroxycampest-4-en-3-one, **14**’s characteristic m/z^+^ signal (MW=414.67Da, [C_28_H_47_O_2_]^+^=415.3) of, i) YYL67 expressing CYP90B1, ii) YYL67 expressing CYP90B1 and CYP90A1, iii) YYL69. **(D)** High resolution mass spectrum of **14**. (E) Quantification of **14** with different promoter combination of CYP90B1 and CYP90A1 expressed in YYL67. All traces and spectrums are representative of at least three biological replicates for each engineered yeast strain and the error bars represent the standard deviation of the replicates.

The conversion efficiency of campesterol to **13** is around 86% (Figure S9). In addition, the activities of an array of sterol C22-hydroxylases were tested in YYL67, including CYP90B1 variants and cholesterol 22-hydroxylases (Table S5). Among all the enzymes, only the analogues of CYP90B1 in *Solanum lycopersicum*, *sl*CYP724B2 and *sl*CYP90B3, can function on campesterol in yeast (Figure S9), as reported before^49^. The membrane bound signaling peptide (~30 amino acids) at the N-terminus was truncated to afford t30CYP90B1, which can convert campesterol to **13**, but with a lower efficiency compared with the full-length protein (Figure S9). This agrees with the previous investigation that the activity of plant cytochrome P450s is retained upon the N-terminus truncation, which can be utilized as an expression strategy in the membrane infrastructure-lacking *E. coli*^50^. Compared with *sl*CYP724B2, *sl*CYP90B3 and t30CYP90B1, CYP90B1 exhibits the highest efficiency towards the synthesis of **13** (Figure S9).

CYP90A1 was first proposed as a C23 hydroxylase^51^, yet subsequent *in vitro* characterization argues that CYP90A1 is more likely a C3 oxidase on BR intermediates with 22-hydroxylated and 22,23-dihydroxylated side chains^52^. Expression of CYP90A1 in YYL67 didn’t lead to the synthesis of either C3-oxidized or C23-hydroxylated product, which agrees with the previous study that campesterol is not a substrate of CYP90A1^52^. Co-expression of CYP90A1 with CYP90B1 in YYL67 led to the synthesis of a new peak of m/z^+^=415.3, which agrees with the putative product 22-hydroxycampest-4-en-3-one, **14**. The LC-MS/MS analysis further confirmed the desaturation of the hydroxyl group on the A-ring (Figure 6D). This functional reconstitution of CYP90A1 and CYP90B1 confirmed the reaction order between these two enzymes. The conversion of **13** to **14** was very efficient, with very little of **13** detected in YYL67 expressing CYP90A1 and CYP90B1 (Figure 6A, 6C), which implies that CYP90B1 may be the rate-limiting step and could be a key to regulate BR biosynthesis in yeast.

We also examined the effect of different promoters on the activities of CYP90A1 and CYP90B1 towards the synthesis of **14**. When CYP90A1 and CYP90B1 are both regulated by strong, constitutive promoters, a slightly higher production of **14** was detected (Figure 6E). We then integrated *CYP90A1* and *CYP90B1* into YYL67 to generate YYL69 (Table S1). Remarkably, the production of **14** was substantially enhanced in comparison to when these two genes were expressed from low-copy plasmids (Table S6), from 0.94 mg/L to 3.63 mg/L, together with an enhanced growth (Figure 6F). This result is surprising and, simultaneously highlights the significance of stable genome integration in pathway reconstruction in *S. cerevisiae*. The establishment of YYL69, the **14**-producing strain, confirms the reaction sequence of CYP90A1 and CYP90B1 in yeast, which is consistent with previous *in planta* and *in vitro* biochemical characterizations. Our work provides a yeast-based platform for the elucidation and engineering of downstream BR biosynthesis.

To further optimize the production of **14** in YYL69, INO2 was overexpressed under the strong, constitutive *GPD* promoter. INO2 was recently found to be able to enlarge the endoplasmic reticulum membrane, where the steroids were synthesized^53^. However, although overexpressing INO2 was very successful in enhancing squalene production, it did not help in the context of sterol production; instead, the titer of **14** dropped ~40% upon the overexpression of INO2 (Figure S10A). Since changing the medium conditions has made substantial differences to the production of campesterol in YYL65, we again explored the effects of various base media and carbon sources on **14** production from YYL69 (Figure 4B). Specifically, YPD resulted in better growth but less production of **14**, in comparison to when cultured in SDM. Across all of the carbon sources examined, glucose exhibited the most promising production of **14**. Further optimization was done through titration between YP and SDM supplemented with 2% (w/v) glucose to locate the optimal balance between **14**-production and cell growth. However, the titer of **14** is proportional to the ratio of SDM (Figure S10B), and the highest production of **14** was still found in SDM with 2% (w/v) glucose. In addition, supplementing 10% (v/v) ethanol in SDM with 2% (w/v) glucose showed ~13% increase in the titer of **14**, with a similar cell density as without ethanol.

## Discussion

Although campesterol production has been previously established in *S. cerevisiae* and *Y. lipolytica* by the function of an animal DHCR7 and the promiscuity of ERG4^22, 23^, such campesterol-producing strain has not been utilized for the synthesis of downstream metabolites, such as BR. In this study, we first established campesterol production using the plant enzymes DWF7, DWF5, and DWF1. Upregulating the mevalonate pathway, a classic strategy to enhance upstream terpene supply, led to more than 10-fold enhancement in campesterol titer to around 40mg/L. However, such high titer of campesterol and its precursors cannot be converted to **13** and the corresponding C24’-methylated sterols upon the introduction of CYP90B1 and SMT2, respectively. Analysis of the campesterol-producing strain YYL56 implies that heterologous sterols in this strain (**6**, campesterol) are only detected in the saponified sample, which indicates the vigorous esterification of these plant specified sterols in yeast (Figure 2A). The inactivation of the sterol acyltransferase genes *are1* and *are2* resulted in the synthesis of **7** and **13**, but in low conversions from their precursors (Figure 1, Figure 3A, 6A). Further inactivation of *erg4*, the paralog of *dwf1*, greatly enhanced the conversion towards the 24’-methylated products such as **7** and β-sitosterol by SMT2, but with a significant growth deficiency. Interestingly, the engineered yeast strains were able to circumvent the growth deficiency through adaptive evolution, with enhanced phytosterol production as well. Although the campesterol titer of the best mutant was only ~7mg/L, less than 1/4 of the highest campesterol titer achieved in this study, the synthesized campesterol is much more accessible by the downstream enzymes, and the C24 stereochemistry is not affected by the presence of ERG4 enzyme. Using this strain, we were able to reconstitute and optimize the early steps of BR biosynthesis, from campesterol to **14**, in yeast with titer of **14** up to ~4mg/L.

Although ERG4 and DWF1 can both function on **6** to afford the synthesis of campesterol, the stereochemistry at C24 can be very different. It seems that introduction of DHCR7 only likely leads to predominant 24(S)-campesterol synthesis. Previous investigations implied that DWF1 from different plants exhibit different stereochemistry, some exclusively catalyze the formation of 24(R)-campesterol, while others result in racemic mixtures^26^. In this study, we used DWF1 from *A. thaliana*. According to previous investigations of AtDWF1^26^ and the high efficiency of ERG4 on **6**, YYL67 likely synthesizes a racemic mixture, while YYL64 synthesizes a mixture with a higher ratio of 24(S)-campesterol. Thus, it is critical to use plant enzymes (DWF1, DWF5, DWF7) to synthesize plant specialized campesterol in yeast, if further downstream pathway is to be constructed.

Through engineering for altered sterol composition, yeast strains were more or less stressed in this study. However, we were surprised to learn the superior stress resistance of *S. cerevisiae*. Apart from the capability to adapt themselves for both enhanced growth and sterol production through simple repeated culture, we also noted that yeast can alter their genetic complement to release the stress. For example, when we re-streaked YYL65 on YPD plate, two phenotypes were detected: there was one colony which grew much faster than remaining majority (Figure S11), which could not produce campesterol and is marked as YYL65-BC. Colony PCR was reperformed on this strain and the previously integrated *dwf5* was no longer detected. In addition, the campesterol production can be rescued by expressing DWF5 from a plasmid in YYL65-BC (Figure S11), which further confirms the loss of function of *dwf5* but not the other integrated genes. On the other hand, we noticed that YYL65 harboring SMT2 showed slightly better growth than YYL65 alone. The enhanced growth in the *dwf5* missing YYL65-BC and SMT2-expressing YYL65 may be attributed to the loss or a decreased level of 24-methylenecholesterol, respectively. We thus hypothesized that 24-methylenecholesterol is toxic to yeast and maybe the major cause of the growth deficiency of YYL67. Additionally, in this study, the ergosterol synthesis was much affected in all the engineered strains, which also implies the flexibility of *S. cerevisiae* to altered sterol composition. Surprisingly, expressing SMT2 in YYL67 led to a slightly decreased titer of β-sitosterol and **7**, compared to YYL65 (Figure 3A). The possible mechanism for the unexpected lower titer of β-sitosterol in YYL67 is not clear, but implies the complexity of sterol metabolism regulation in *S. cerevisiae*.

Similar to many previously reported metabolic engineering efforts, different medium conditions can cause distinct effects on the target compound production. YYL65 exhibits a significant growth deficiency (Figure 5B) that cannot be used to express cytochrome P450s, and was not able to grow in a number of medium conditions. YYL65 grows slightly better in glucose, yet exhibits a higher campesterol titer in sucrose. This agrees with previous investigations that ergosterol production tends to be higher when using dihexoses (e.g. sucrose and maltose) as carbon sources in yeast, compared to monosaccharides (e.g. glucose)^54^. However, such a trend was diminished in the adapted strain YYL67 (Figure S6B). Upon the introduction of plant cytochrome P450s, however, glucose became a more favorable carbon source in respect of both sterol synthesis and growth (Figure 4B). Unlike previous studies that addition of glycerol or trehalose helped the noscapine production in *S. cereivsiae*^55^, addition of glycerol did not help the production of **14** and YYL69 cannot grow in the presence of trehalose (Figure 4B). Addition of ethanol did enhance the titer of **14** but the effect was subtle, which might be due to the positive effect of ethanol on acetyl-coA supply and sterol biosynthesis as discussed previously. The distinct preference in carbon sources between YYL65, YYL67, and YYL69 may shed light on potential engineering strategies for phytosterol-production and plant cytochrome P450 reconstitution in yeast.

Through reconstituting the biosynthesis of β-sitosterol and **14**, we demonstrated the critical bottlenecks in converting phytosterol to downstream products in yeast - crosstalk between heterologous and yeast endogenous sterol metabolism, and the growth burden caused by altered sterol composition in yeast. We believe that these challenges are generally true in the reconstitutions of other phytosterol-derived pathways in yeast. Thus, this work has broad implications on engineering strategies to establish phytosteroid-producing yeast strains in general. In addition, this work highlights the potential of using baker’s yeast to elucidate and engineer the biosynthesis of various steroids, such as steroidal alkaloids and withanolides. The establishment of a **14**-producing strain will provide a platform to elucidate the biosynthesis of BRs. Thus, *S. cerevisiae* is an optimal host for phytosterol pathway elucidation and reconstitution. In addition, considering the growth deficiency in YYL67 and YYL69, non-conventional yeast strains with more robust growth or higher lipid production, such as *Y. lipolytica*, should also be considered for the manufacturing of phytosteroids once the biosynthesis is fully elucidated and reconstituted in *S. cerevisiae*.

## Materials and Methods

### Materials and culture conditions

Chemicals: campesterol (~98%) and ergosterol (≥75%) are obtained from Sigma-Aldrich. β-Sitosterol (~65%) and stigmasterol (~95%) are purchased from Fisher Scientific. All engineered yeast strains in this work are listed in Table S1 and constructed in a haploid CENPK2.1D background (*MATα; his3D;1 leu2-3_112; ura3-52; trp1-289; MAL2-8c; SUC2*). Yeast strains were cultured at 30 °C in complex yeast extract peptone dextrose (YPD, all components from BD Diagnostics) medium or synthetic defined medium (SDM) containing yeast nitrogen base (YNB) (BD Diagnostics), ammonium sulfate (Fisher Scientific), 2% (w/v) glucose unless specified and the appropriate dropout (Takara Bio) solution for selection. 200 mg/L G418 sulfate (Calbiochem) or 200 mg/L Hygromycin B (Life Technologies) were used in YPD medium for selection.

### General technique for DNA manipulation

PCR reactions were performed with Expand high Fidelity system (Sigma-Aldrich), Phusion DNA Polymerase (NEB), Q5 High-Fidelity DNA Polymerase (New England Biolabs) and Taq Polymerase (NEB) according to manufacturer’s protocols. PCR products were purified by Zymoclean Gel DNA Recovery Kit (Zymo Research). Plasmids were prepared with Econospin columns (Epoch Life Science) according to manufacturer’s protocols. All DNA constructs were confirmed through DNA sequencing by Source Bioscience Inc and Retrogen sequencing Inc. BP Clonase II Enzyme Mix, Gateway pDONR221 Vector and LR Clonase II Enzyme Mix (Life Technologies) and the *S. cerevisiae* Advanced Gateway Destination Vector Kit^56^ (Addgene) were used to perform Gateway Cloning. Gibson one-pot, isothermal DNA assembly^57^ was conducted at the scale of 10 μL by incubating T5 exonuclease (NEB), Phusion polymerase (NEB), Taq ligase (NEB) and 50 ng of each DNA fragment at 50 °C for 1 h to assemble multiple DNA fragments. Yeast strains are constructed through homologous recombination and DNA assembly^58^. *ATR1, DWF1, DWF5, DWF7, SMT2* were amplified from *A. thaliana* cDNA (kindly provided by Prof. Patricia Springer). *DHCR7* from Xenopus laevis, *CYP90A1, CYP90B1, CYP90D1* and *DET2* from *A. tahliana* were *S.cerevisiae* codon optimized and synthesized from TWIST Bioscience Inc. Plasmids used in this study are in Table S5. Sequences of genes used in this work are listed in Table S6.

### Culture and fermentation conditions

All yeast strains were grown in 500 μL media in 96-well plates (BD falcon) covered with AeraSeal film (Excel Scientific), shaking at 250 r.p.m.. For all the engineered yeast metabolites analysis, yeast strains were first cultured overnight in 500 μL SDM with 2% (w/v) glucose. Appropriate volume of the overnight seed culture was inoculated in fresh 500 μL SDM to make the OD600 around 0.1 and incubated at 30 °C for 72 h before metabolite analysis of the yeast pellets.

### Analysis of phytosterol and intermediates in BR pathway

All yeast metabolites of interests in this study were all collected from yeast pellets. For the strains producing steryl esters, 100 μL 60% (w/v) KOH and 100 μL ethanol was added to each sample, then incubated at 86 °C for 1 to 2 hours for saponification. The mixture was cooled down to room temperature and then 400 μL petroleum ether was added and vortexed thoroughly for 15 min. Organic phases were removed to new tubes and dried in vacufuge (Eppendorf). For Δare1Δare2 double mutants, 100 μl of dimethylformamide (DMF) was added to each sample. Then 400 μl of petroleum ether was add and vortex to mix. The petroleum ether phase was taken to new tubes and dried in the vacufuge. For the intermediates in BR synthetic pathway, yeast pellets from 1 ml culture were collected and mixed with 100 μl of DMF and 900 μl acetone. The mixture was vortexed for 15 min and spun at 13.3 krpm for 10 min. The liquid phases were taken to new tubes and dried in the vacufuge. All dried samples were eluted in ethanol and run through LC-MS.

LC-MS used here is Shimadzu serial 2020. The column used in analysis is from Agilent HPLC column Poroshell 120, EC-C18, 3.0 x 100 mm, 2.7 μm. On Shimadzu LC-MS 2020, phytosterols were separated on isometric acetonitrile: methanol (80: 20, v/v, 0.1% formic acid) over 15 min with 0.6 ml/min flow rate at 25 °C. The intermediate **13**, **14**, **15** and **16** on BR pathway were separated under the linear gradient started from 80% (v/v) methanol in water (1% formic acid) to 100% (v/v) methanol over 8 min and then stay at 100% (v/v) methanol for 12 min at the flow rate of 0.5 ml/min. The mass spectrometer was operated in positive ionization mode. Data was acquired by the electrospray ionization mode. Characteristic m/z+ of the compounds was set in SIM mode as described in figure captions.

### Quantification of campesterol, sitosterol and intermediates on BR synthetic pathway

For the quantification of campesterol, calibration curve of mass abundance to concentration was made with the authentic campesterol standard solution. Mass abundance was represented by the peak area at SIM m/z^+^=383.3 mode, which is automatically calculated by Labsolution of Shimadzu LC-MS 2020.For the intermediates **13**, **14**, **15** and **16** on BR pathway, the authentic standards are not available. The concentrations of them were reported as equivalents of campesterol using the standard curve of campesterol and the relative abundance of characteristic ion signals of the corresponding compounds: SIM m/z^+^=399.3, 415.3, 439.3, 455.4, respectively.

### Adaptive evolution

Seed culture of yeast strain YYL65 was started with a single colony from an agar plate and grown to stationary phase. Then the culture was back-diluted in 2mL synthetic complex SDM with 2% (w/v) sucrose, starting at OD_600_~0.1 to early stationary phase (OD600 ~6.0). Then appropriate amount of the culture was used to inoculate 2mL fresh synthetic complete SDM with 2% (w/v) sucrose, with the starting OD_600_~0.1. This procedure was repeated for more than 10 rounds. Then 10uL culture from the last batch of culture was plated on synthetic complex SDM agar plate. 10 colonies were cultured in 1mL SDM liquid medium at 30 °C for 3 days, which are subjected to LC-MS analysis for campesterol production.

### Genome sequencing

Genomic DNA was extracted with VWR Life Science Yeast Genomic DNA Purification Kit, following the standard protocol from the vendor. The sequencing was done by Novogen Corporation. The chromatogram files generated by NGS platforms (like Illumina HiSeq TM 2000, MiSeq) are transformed by CASAVA Base Calling into sequencing reads, which are called Raw data or Raw reads. Both the sequenced reads and quality score information would be contained in FASTQ files. Raw data is filtered off the reads containing adapter and low-quality reads to obtain clean data for subsequent analysis. The resequencing analysis is based on reads mapping to a common reference sequence by BWA software. SAMTOOLS is used to detect SNP and InDel in functional genomics and get the mutation statistics.

### LC-MS/MS analysis of intermediates on BR pathway

Structure of intermediates **13**, **14**, **15** and **16** on BR pathway were identified by Waters Synapt G2-Si Q-TOF system. Yeast sample preparation and LC chromatograph were the same as described in “Analysis of phytosterol and intermediates in BR pathway” session. The mass spectrometer was operated in positive ionization mode. MS/MS fragmentation data was obtained using Collision Induced Dissociation (CID) fragmentation, with collision voltage at 25 V.

### Yeast growth monitoring

Yeast growth was monitored with OD600 measurement. Yeast grew in SDM medium in 96-well plates as described in “Culture and fermentation conditions” session. The yeast culture was collected into microplate (from Coring) at certain time intervals after induction, and diluted with blank SDM medium if necessary. The data was obtained by Synergy HTX Multi-Mode Microplate reader from Bio Tek. The reading from microplate reader was calibrated with spectrophotometer and then converted to OD600.

## Acknowledgements

We thank the Metabolomics Core Facility at UC Riverside and Dr. Jay Kirkwood for instrument access, training, and data analysis; Shibo Wang for the valuable suggestions and discussions on genome sequencing analysis; Dr. Sheng Wu, Tiffany Chiu, and Anqi Zhou for valuable feedback and discussion in the preparation of the manuscript. This work was supported by the National Science Foundation (grant to Y.L., 1748695) and Cancer Research Coordinating Committee Research Award (grant to Y.L., CRN-20-634571).

## Author contributions

S.X. and Y.L. conceived of the project, designed the experiments, analyzed the results and wrote the manuscript. S.X. and C.C. performed the experiments.

