## Supplemental Information for "Engineering of Phytosterol-Producing Yeast Platforms for Functional Reconstitution of Downstream Biosynthetic Pathways"

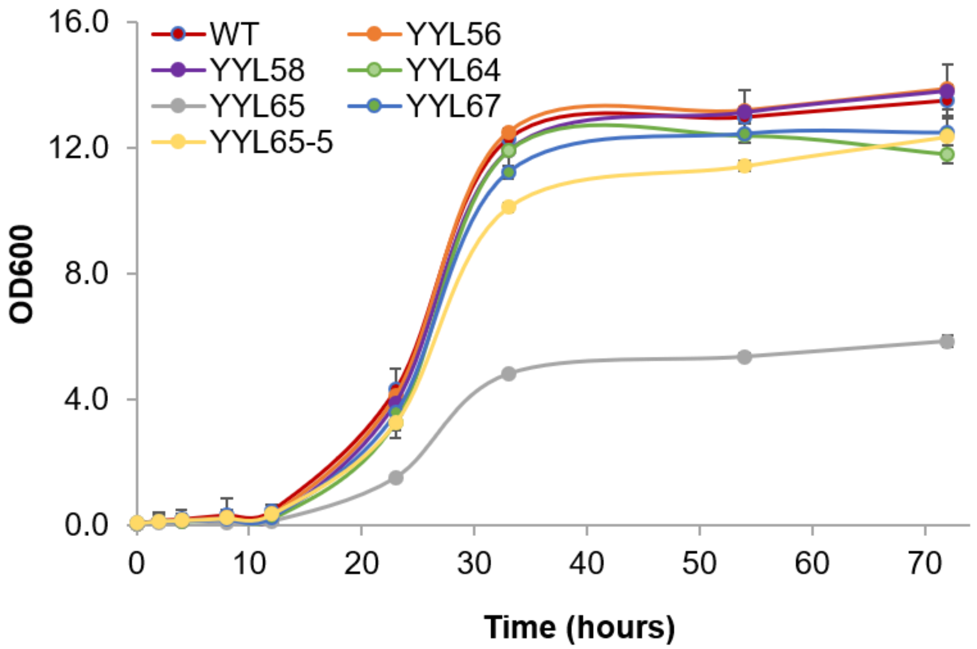


**Figure S1.** Growth curve of strains engineered for campesterol production. The yeast strains were cultured in synthetic complete SDM medium supplemented with 2% (w/v) glucose at 30°C for 72 hours. Bars represent mean values of three biological replicates, and the error bars represent the standard deviation of the replicates.

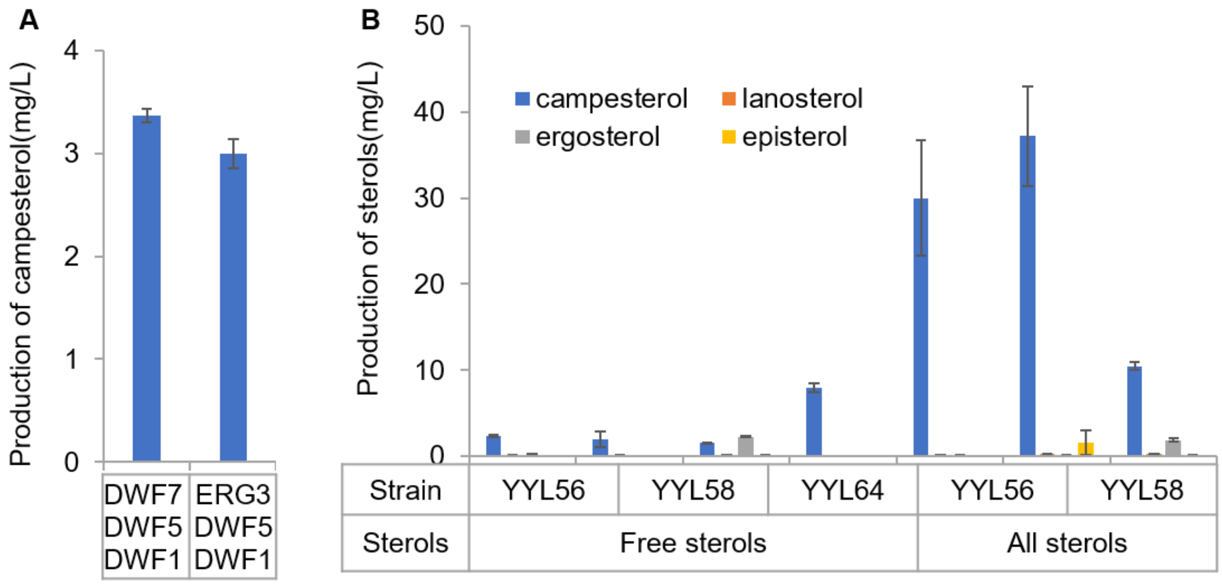


**Figure S2.** Sterol profiling of YYL56, YYL58 and YYL64. Lanosterol, episterol, ergosterol and campesterol was quantified in free sterols and total sterols including sterol esters, by using unsaponified and saponified extraction methods. Bars represent mean values of three biological replicates, and the error bars represent the standard deviation of the replicates. The quantification of each sterol was measured by comparing the integrated peak area to a standard curve of the authentic standards of the corresponding sterols.

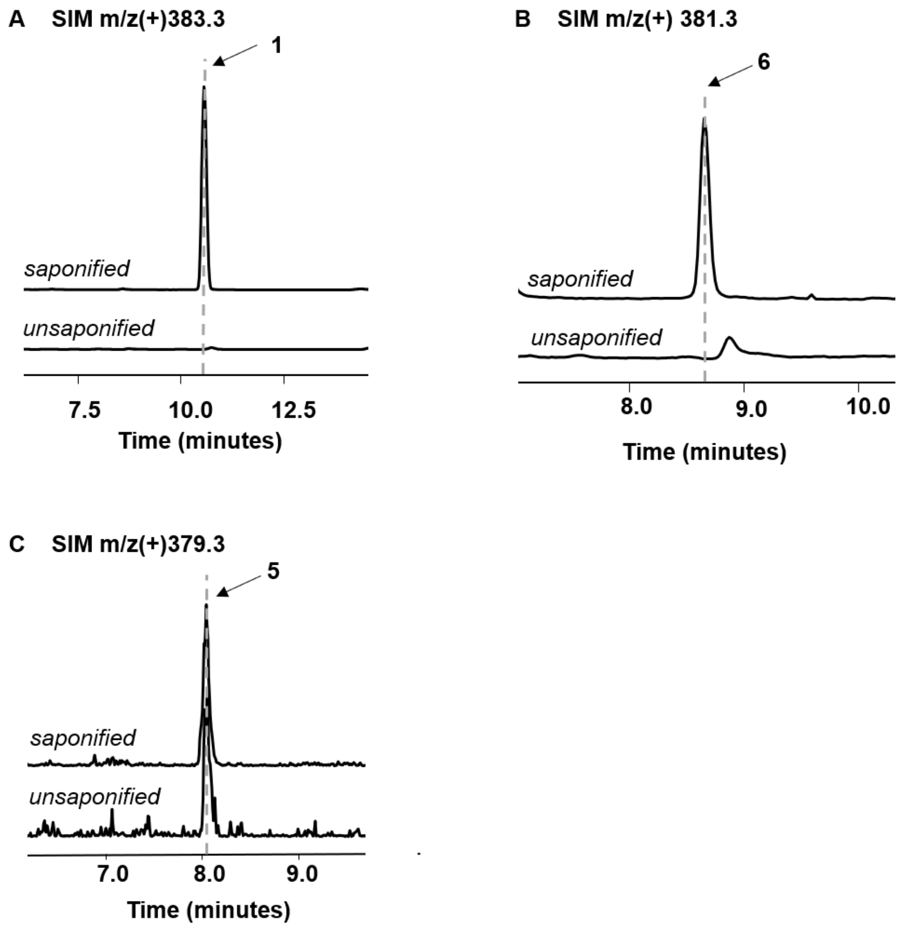


**Figure S3.** LCMS traces of campesterol and **6** in saponified and unsaponified extraction of YYL58. (A) EIC of campesterol characteristic m/z+ =383.3 of the saponified and unsaponified sterol extract of YYL58. (B) EIC of compound **6** characteristic m/z+ =381.3 of the saponified and unsaponified sterol extract of YYL58. (C) EIC of compound **5** characteristic m/z+ =379.3 of the saponified and unsaponified sterol extract of YYL60. All traces are representative of at least three biological replicates for each engineered yeast strain.

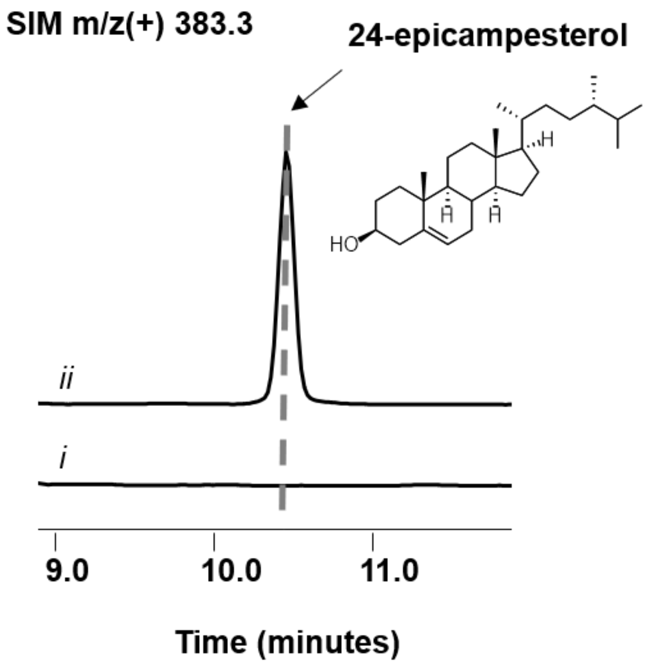


**Figure S4.** LC-MS analysis of the function of DWF5 in yeast strains with or without *erg4*. SIM EIC of characteristic campesterol or 24-epicampesterol characteristic m/z$^+$=383.3 of i) YYL66 expressing DWF5, ii) YYL63 expressing DWF5. Traces are representative of at least three biological replicates for each engineered yeast strain.

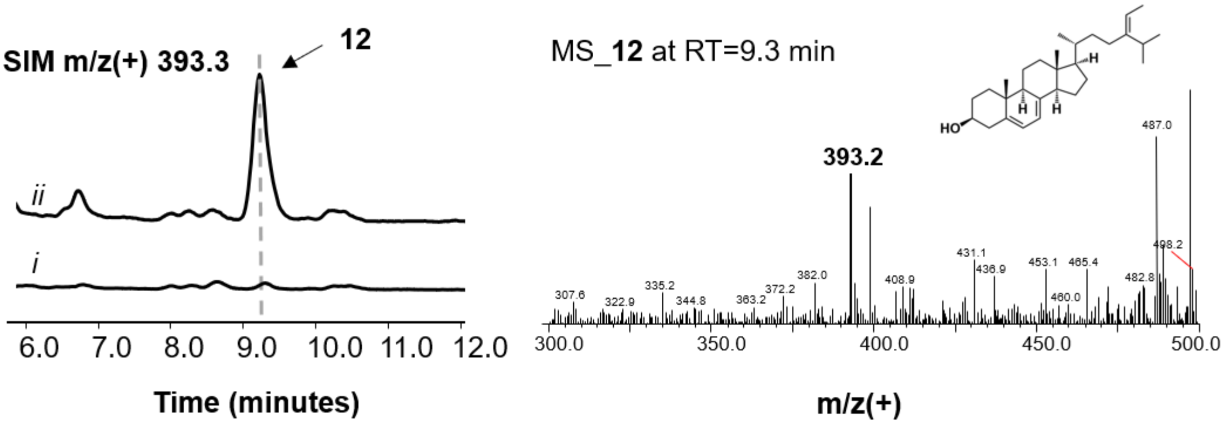


**Figure S5.** EIC SIM using Δ5,7-Avenasterol, **12**, characteristic m/z$^+$ signal (MW=410.69, $[C_{29}H_{45}]^+$=393.3) in i) **5**-accumulated strain YYL60, ii) YYL60 harboring SMT2. The mass spectrum of the peak at 9.3 minutes is shown on the right. The traces and spectrums are representative of at least three biological replicates for each engineered yeast strain.

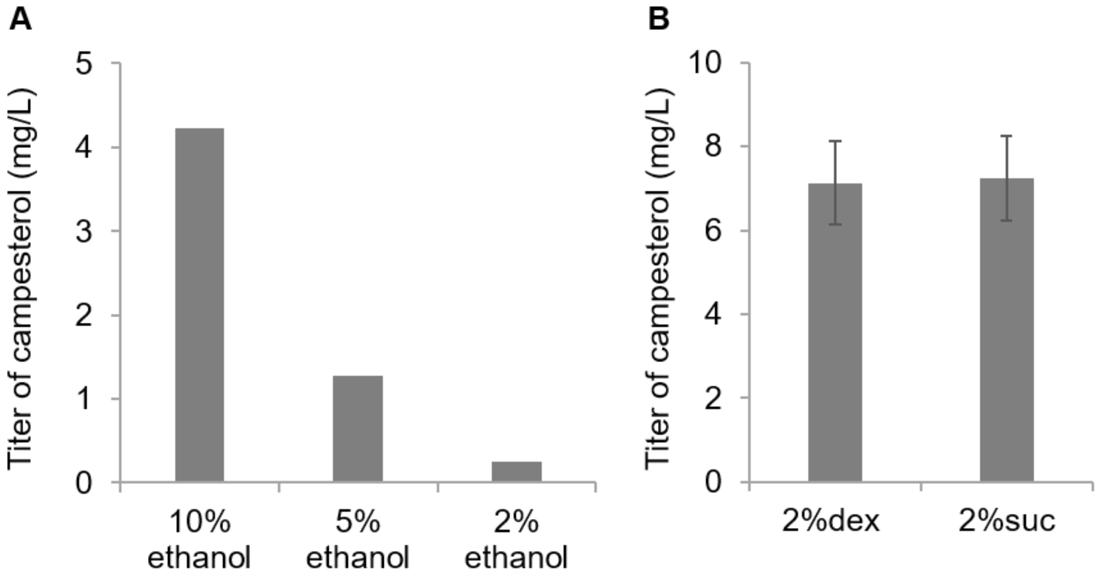


**Figure S6**. (A) ethanol titration of YYL65 for medium optimization. YYL65 was cultured in SDM medium with 2% sucrose, adding 10%, 5% and 2% ethanol. (B) use 2% glucose (dex) and sucrose (suc) as carbon source in culturing YYL67 in SDM. Bars represent mean values of three biological replicates, and the error bars represent the standard deviation of the replicates.

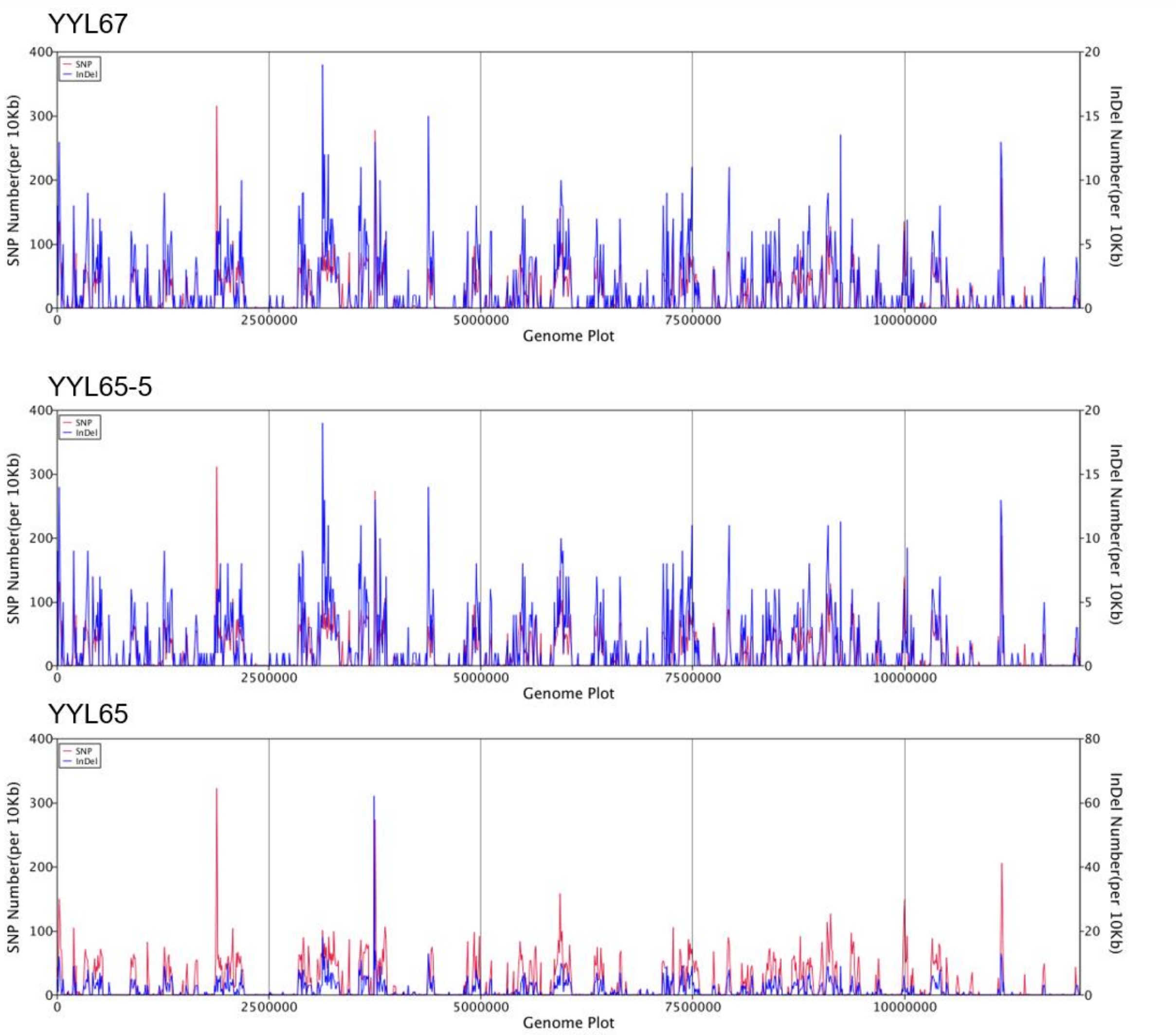


**Figure S7.** SNP/InDel distribution through the whole genome of strain YYL67, YYL65-5 and YYL65. Numbers of SNPs are represented by red line. Numbers of InDels are represented by blue line. The genome information of *S. cerevisiae* S288c is utilized as the reference genome here for alignment and annotation.

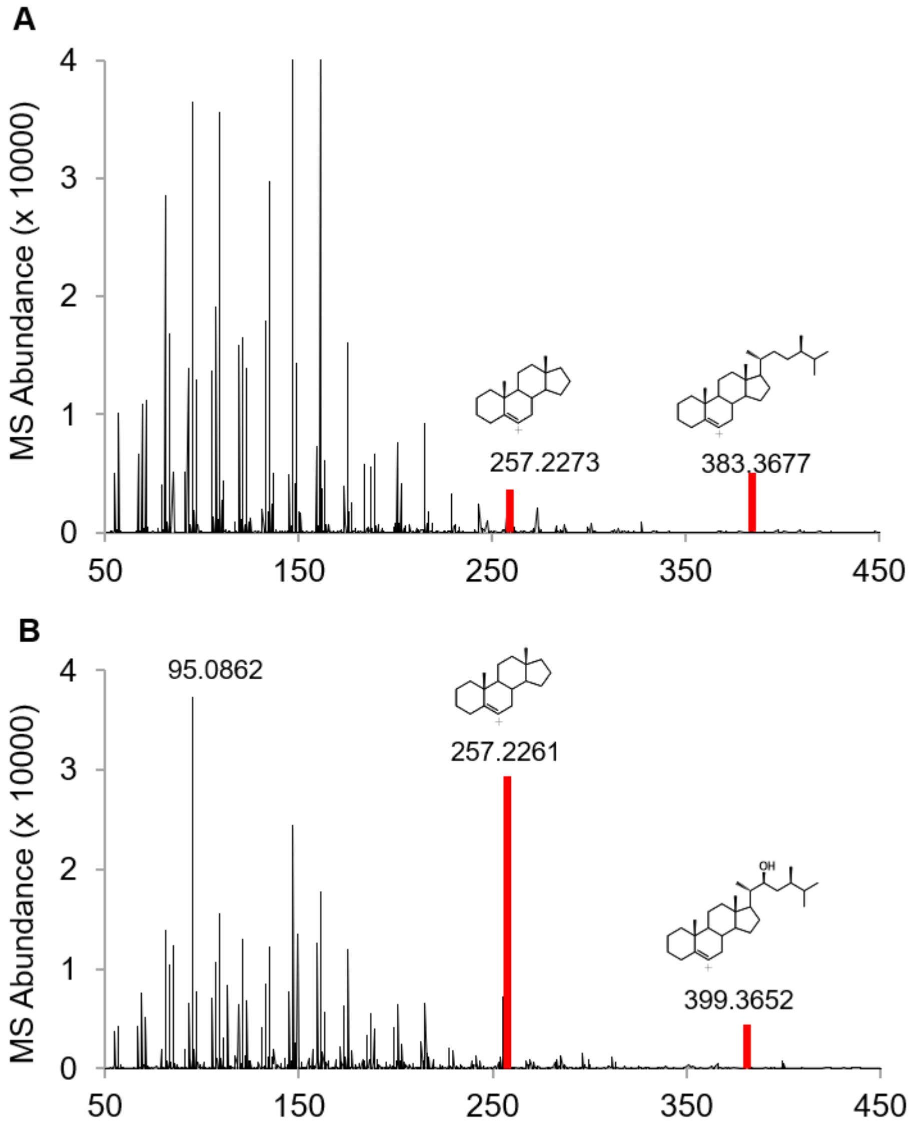


**Figure S8.** Mass spectrum of **(A)** campesterol, **(B) 13**. The spectrums are representative of at least three biological replicates for each engineered yeast strain.

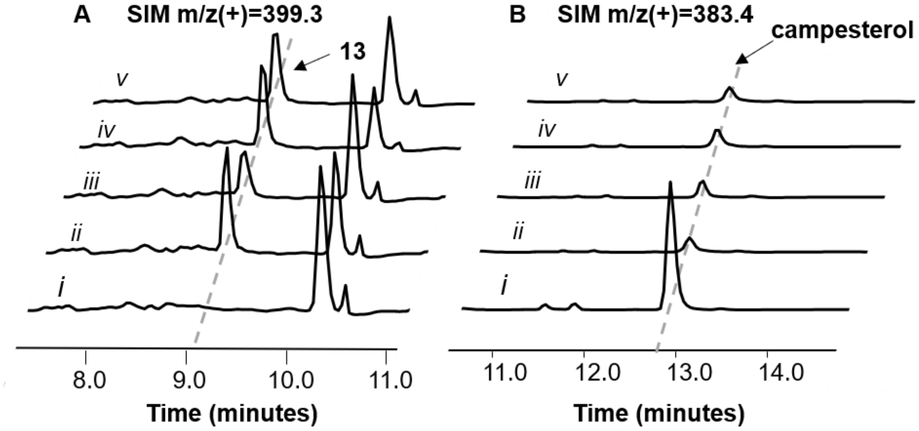


**Figure S9.** Characterization the activities of CYP90B1 variants and mutants. **(A)** EIC SIM using 22-hydroxycampesterol, **13**, characteristic m/z$^+$ signal (MW=416.69, $[C29H45]^+$=399.4) in YYL67 expressing i) empty vector, ii) CYP90B1, iii) CYP724B2, iv) CYP90B3, v) t30CYP90B1. **(B)** EIC SIM using campesterol's characteristic m/z$^+$ signal (MW=400.69, $[C29H45]^+$=383.4) in YYL67 expressing i) empty vector, ii) CYP90B1, iii) CYP724B2, iv) CYP90B3, v)t30CYP90B1. YYL67 harboring plasmids with CYP90B1 variants and mutants (pYL655-pYL663) were grown in selective SDM supplemented with 2% glucose at 30°C for 72 hours before analysis. Only the enzymes that converted campesterol were presented, and the full list of CYP90B1 variants tested are listed in Table S3.

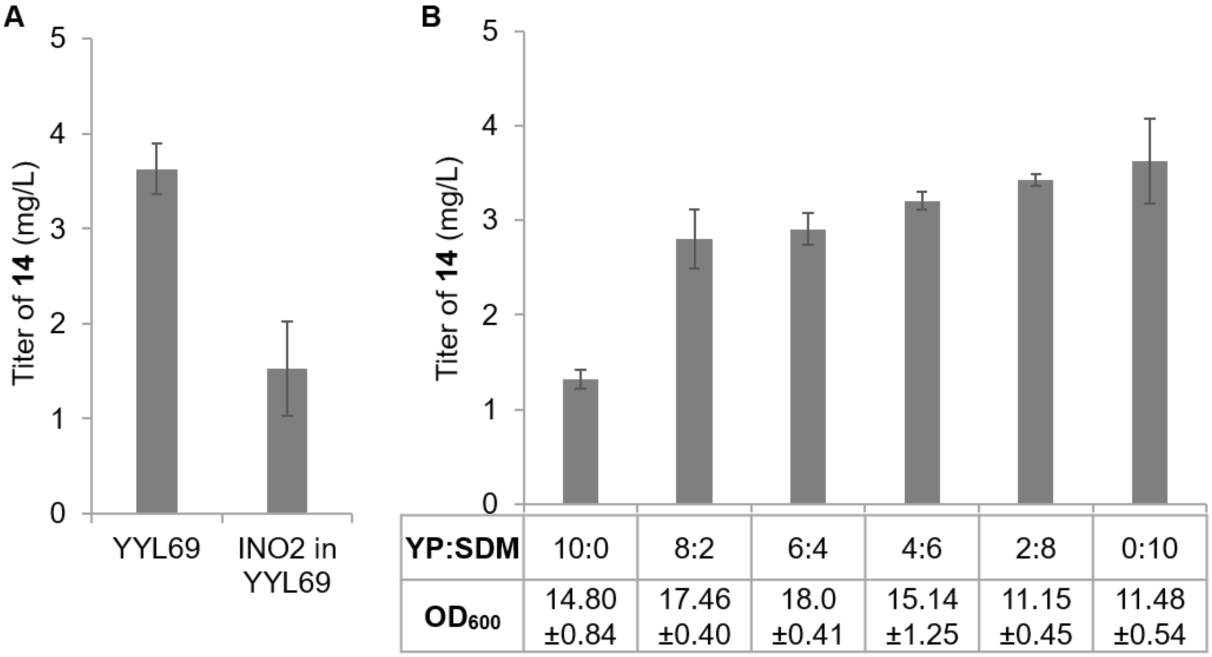


| YP:SDM | 10:0 | 8:2 | 6:4 | 4:6 | 2:8 | 0:10 |
|---|---|---|---|---|---|---|
| $OD_{600}$ | 14.80 ±0.84 | 17.46 ±0.40 | 18.0 ±0.41 | 15.14 ±1.25 | 11.15 ±0.45 | 11.48 ±0.54 |

**Figure S10.** Optimization of the production of **14**. **(A)**YYL69 harboring plasmid $P_{GPD}$-INO2 and YYL69. Yeast strains were grown in synthetic complete SDM medium at 30°C for 72 hours. **(B)** Titration of YP to SDM ratio with 2% (w/v) glucose. Bars represent mean values of three biological replicates, and the error bars represent the standard deviation of the replicates.

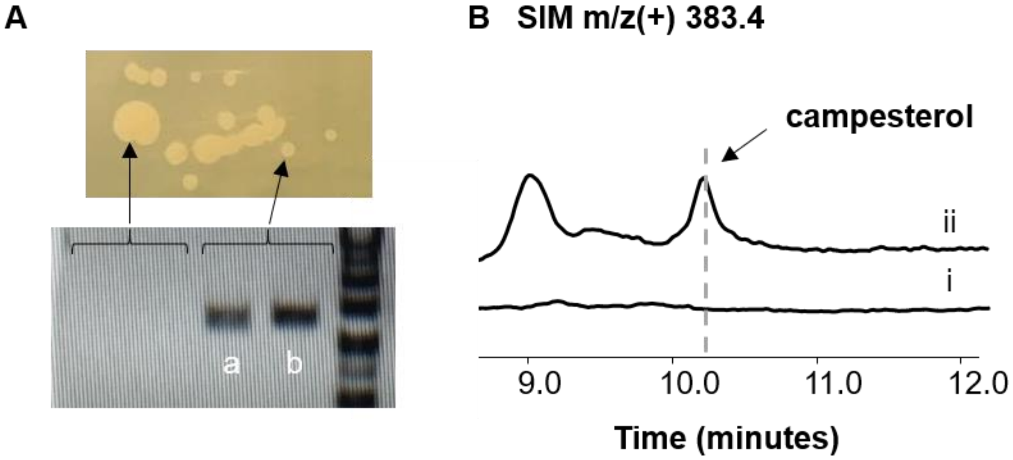


**Figure S11**. Identification of DWF5 in YYL65 big colony (BC) mutant. (A) colony PCR of DWF5 integration boundaries in YYL65 and YYL65-BC mutant indicating the loss of *dwf5*. a and b indicated the left and right boundaries of *dwf5*. (B) EIC SIM using campesterol's characteristic m/z$^+$ signal (MW=400.69, $[C29H45]^+$=383.4) in i) YYL65-BC and ii) YYL65-BC expressing DWF5.

**Table S1.** Genotypes of yeast strains utilized in this study.

| **Strain** | **Genotype** |
|---|---|
| YYL55 | *leu2Δ::$P_{PYK1}$-dwf1-$T_{MFA1}$, $P_{TPI1}$-atr1-$T_{STE2}$, $P_{PGK1}$-dwf5-$T_{PHO5}$, kanmx, $P_{GPD}$-dwf7-$T_{CYC1}$* |
| YYL56 | *leu2Δ::$P_{PYK1}$-dwf1-$T_{MFA1}$, $P_{TPI1}$-atr1-$T_{STE2}$, $P_{PGK1}$-dwf5-$T_{PHO5}$, $P_{GPD}$-dwf7-$T_{CYC1}$*<br>*ybl059wΔ::$P_{PGK1}$-erg12-$T_{PHO5}$, $P_{TEF1}$-erg10-$T_{CYC1}$, $P_{PYK1}$-thmg1-$T_{MFA1}$, $P_{TPI1}$-erg13-$T_{STE2}$* |
| YYL57 | *leu2Δ::$P_{PYK1}$-dwf1-$T_{MFA1}$, $P_{TPI1}$-atr1-$T_{STE2}$, $P_{PGK1}$-dwf5-$T_{PHO5}$, $P_{GPD}$-dwf7-$T_{CYC1}$*<br>*ybl059wΔ::$P_{PGK1}$-erg12-$T_{PHO5}$, $P_{TEF1}$-erg10-$T_{CYC1}$, $P_{PYK1}$-thmg1-$T_{MFA1}$, $P_{TPI1}$-erg13-$T_{STE2}$*<br>*ybr197cΔ::$P_{TPI1}$-erg8-$T_{STE2}$, $P_{TEF1}$-erg19-$T_{CYC1}$, $P_{GPD}$-idi1-$T_{CYC1}$, $P_{PYK1}$-thmg1-$T_{MFA1}$* |
| YYL58 | *leu2Δ::$P_{PYK1}$-dwf1-$T_{MFA1}$, $P_{TPI1}$-atr1-$T_{STE2}$, $P_{PGK1}$-dwf5-$T_{PHO5}$, $P_{GPD}$-dwf7-$T_{CYC1}$*<br>*ybl059wΔ::$P_{PGK1}$-erg12-$T_{PHO5}$, $P_{TEF1}$-erg10-$T_{CYC1}$, $P_{PYK1}$-thmg1-$T_{MFA1}$, $P_{TPI1}$-erg13-$T_{STE2}$*<br>*ybr197cΔ::$P_{TPI1}$-erg8-$T_{STE2}$, $P_{TEF1}$-erg19-$T_{CYC1}$ $P_{GPD}$-idi1-$T_{CYC1}$, $P_{PYK1}$-thmg1-$T_{MFA1}$*<br>*ymr206wΔ::$P_{TEF1}$-erg20-$T_{CYC1}$, $P_{GPD1}$-upc2 -$T_{ADH1}$, $P_{PYK1}$-thmg1-$T_{MFA1}$, $P_{PGK1}$-erg7-$T_{PHO5}$* |
| YYL60 | *erg5Δ::dhcr7* |
| YYL63 | *are1Δ, are2Δ* |
| YYL64 | *leu2Δ::$P_{PYK1}$-dwf1-$T_{MFA1}$, $P_{TPI1}$-atr1-$T_{STE2}$, $P_{PGK1}$-dwf5-$T_{PHO5}$, $P_{GPD}$-dwf7-$T_{CYC1}$*<br>*ybl059wΔ::$P_{PGK1}$-erg12-$T_{PHO5}$, $P_{TEF1}$-erg10-$T_{CYC1}$, $P_{PYK1}$-thmg1-$T_{MFA1}$, $P_{TPI1}$-erg13-$T_{STE2}$*<br>*are1Δ, are2Δ* |
| YYL65 | *leu2Δ::$P_{PYK1}$-dwf1-$T_{MFA1}$, $P_{TPI1}$-atr1-$T_{STE2}$, $P_{PGK1}$-dwf5-$T_{PHO5}$, $P_{GPD}$-dwf7-$T_{CYC1}$*<br>*ybl059wΔ::$P_{PGK1}$-erg12-$T_{PHO5}$, $P_{TEF1}$-erg10-$T_{CYC1}$, $P_{PYK1}$-thmg1-$T_{MFA1}$, $P_{TPI1}$-erg13-$T_{STE2}$*<br>*are1Δ, are2Δ*<br>*erg4Δ::ura3* |
| YYL66 | *are1Δ, are2Δ*<br>*erg4Δ::ura3* |
| YYL67 | *leu2Δ::$P_{PYK1}$-dwf1-$T_{MFA1}$, $P_{TPI1}$-atr1-$T_{STE2}$, $P_{PGK1}$-dwf5-$T_{PHO5}$, $P_{GPD}$-dwf7-$T_{CYC1}$*<br>*ybl059wΔ::$P_{PGK1}$-erg12-$T_{PHO5}$, $P_{TEF1}$-erg10-$T_{CYC1}$, $P_{PYK1}$-thmg1-$T_{MFA1}$, $P_{TPI1}$-erg13-$T_{STE2}$*<br>*are1Δ, are2Δ*<br>*erg4Δ* |
